# Watkins wheat landraces: a treasure of stripe rust resistance alleles identified using multi-model association analyses

**DOI:** 10.64898/2026.03.11.711137

**Authors:** Jasneet Singh, Muhammad Jawad Akbar Awan, Naveen Kumar, Samuel Holden, Rajdeep Khangura, Gurcharn S. Brar

## Abstract

Wheat stripe rust, caused by *Puccinia striiformis* f. sp. *tritici* (*Pst*), remains a major global constraint to wheat production. Rapid pathogen evolution, exemplified by the recent breakdown of *Yr15* in Europe, underscores the need to identify diverse and durable resistance loci. The A.E. Watkins landrace collection represents a globally diverse pre-breeding resource with substantial untapped variation for stripe rust resistance. In this study, 297 Watkins landraces were evaluated against six diverse *Pst* isolates (representing six races and three North American lineages) and subjected to genome-wide association analysis using high-density whole-genome resequencing data. Continuous phenotypic variation was observed across isolates, with several accessions displaying stable resistance across all lineages. A total of 87 QTLs were identified across all 21 wheat chromosomes. Ten loci co-localized with designated or cloned *Yr* genes, including *Yr84*, *Yr85*, *Yrq1*, *Yr71*, *Yr60*, *Yr62*, *Yr50*, *Yr68*, *Yr34*, and *Lr34/Yr18/Sr57*. An additional 34 loci overlapped previously reported stripe rust QTL, whereas the majority did not coincide with known loci, suggesting potential novel resistance regions. Eighteen QTLs were supported by multiple isolates, and fourteen showed supports across statistical models, indicating robust genomic signals. Several Watkins accessions carried favorable alleles that co-localized with multiple *Yr*-aligned loci, identifying promising donor candidates for validation and pre-breeding.

**Key Message:** Genome-wide association mapping of 297 Watkins wheat landraces across diverse stripe rust races & genetic lineages identified 87 *QTL*, including 10 formally designated *Yr* genes and 46 novel loci, highlighting Watkins landraces as valuable pre-breeding donors for novel all-stage stripe rust resistance.

## Introduction

Wheat stripe or yellow rust (*Yr*), caused by the obligate biotrophic fungus *Puccinia striiformis* f. sp. *tritici* (*Pst*), remains one of the most destructive diseases of wheat worldwide (Chen 2020). The pathogen evolves rapidly through mutation, clonal expansion, occasional recombination, and long-distance wind dispersal, enabling frequent shifts in virulence that undermine deployed resistance genes (Ali et al. 2014; McDonald and Linde 2002; Holden et al. 2025). The recent breakdown of the widely deployed resistance gene *Yr15* in the UK and parts of Europe, following the emergence of a *Yr15*-virulent *Pst* race in 2025, underscores the vulnerability of single-gene resistance and reinforces the need to continuously identify and deploy diverse resistance loci for durable stripe rust control (Davis et al. 2025).

Global wheat breeding for high yield and agronomic uniformity has progressively reduced the genetic diversity among modern cultivars due to recurrent use of related parental lines and strong directional selection (Niu et al. 2023; Semagn et al. 2021). This restricted genetic base limits the reservoir of available resistance alleles and increases vulnerability to emerging pathogen races. Expanding the gene pool through systematic exploration of diverse and underutilized germplasm can offer novel alleles for multiple traits of economic importance (particularly disease resistance) and enrich genetic diversity in the elite gene pool. The A.E. Watkins bread wheat collection comprises 827 hexaploid landrace accessions assembled in the 1920s and 1930s from across 32 countries, capturing a global snapshot of wheat diversity prior to modern breeding (Anonymous 2024; Wingen et al. 2017).

Our program (Cereal Breeding Lab) started working on a smaller subset of Watkins landrace collection in 2019 with an aim to identify disease resistance and develop pre-breeding germplasm. The focus for last few years has been on mining resistance to stripe rust and Fusarium head blight (FHB), two of the five priority diseases of wheat in Canada (Brar et al. 2019). The Watkins collection has proved to be a reliable reservoir of stripe rust resistance. Multiple independent studies have identified novel *Yr* genes from some selected Watkin accessions, including *Yr47* (Bansal et al. 2011), *Yr51* (Randhawa et al. 2014), *Yr57* (Randhawa et al. 2015), *Yr60* (Bariana et al. 2014), *Yr63* (Mackenzie et al. 2023), *Yr72* (Chhetri et al. 2023), *Yr80* (Nsabiyera et al. 2018), *Yr81* (Gessese et al. 2019), and *Yr82* (Pakeerathan et al. 2019). These genes span multiple chromosomes and represent both seedling and adult-plant resistance types. Advancing this resource, whole-genome re-sequencing of all 827 Watkins accessions alongside 208 globally diverse modern cultivars (≈12.73× reported depth for Watkins) has delivered a high-resolution variant and haplotype map and resolved seven ancestral groups within Watkins panel (Cheng et al. 2024). Modern wheat was shown to derive predominantly from only two of these groups, implying that the remaining groups harbour underutilized alleles and long-range haplotypes absent from elite germplasm (Cheng et al. 2024).

Here, we leveraged the genomic resource developed by Cheng et al. (2024) and conducted a genome-wide association study (GWAS) on a subset of 297 Watkins accessions evaluated against six diverse *Pst* isolates representing the most predominant *Pst* races and genetic lineages in North America. Our study reports genomic regions corresponding to several previously formally designated *Yr* genes while also identifying novel resistance loci not overlapping with any known genes. Our study reports a comprehensive list of *Yr* genes/QTLs carried by a small subset of Watkins landrace panel. Significant *Yr*-associated loci emerging from this analysis will be prioritized for validation and conversion into breeder-friendly markers to support pre-breeding and introgression.

## Materials and methods

### Plant material

A subset of 297 hexaploid wheat landraces from the A.E. Watkins collection was obtained from the John Innes Centre, UK, in 2019 (**Table S1**). These landraces were used for all phenotypic and genotypic analyses conducted in this study.

### Stripe rust phenotyping

The panel of 297 Watkins landraces was planted in 72-cell plastic trays with one plant per cell. The panel was grown in two replications under controlled growth chamber conditions. For phenotyping, six *Pst* isolates representing diverse genetic lineages were used **(Table 1)**. Twelve-day-old seedlings were inoculated with a urediniospore suspension of each isolate individually. The isolates used in this study included W001 and W034 (genetic lineage PstS1), W031 and W057 (PstS18), W056 (PstPr), and T022 (unknown lineage) (**Table 1**) (Brar and Kutcher 2016). Urediniospores were suspended in light mineral oil (Novec 7100™) and sprayed evenly onto the seedlings using an air compressor. After inoculation, plants were incubated in a dark dew chamber at 10°C for 24 h to facilitate infection and were subsequently transferred to a growth chamber maintained at approximately 16°C with a 16 h light/8 h dark photoperiod. Infection types (ITs) were recorded 14–16 days after inoculation using the standard 0–9 scale (Line and Qayoum 1992), where IT of 0-3 was considered resistant, 4-6 as partially resistant, and IT of 7-9 as susceptible.

**Table 1.** *Pst* isolates used for stripe rust phenotyping of the Watkins landrace panel and their genetic lineages.

| Isolate name | Genetic lineage |
| --- | --- |
| W001 | PstS1 |
| W031 | PstS18 |
| W034 | PstS1 |
| W056 | PstPr |
| W057 | PstS18 |
| T022 | Unknown |

### Genotypic data

Whole-genome variant call format (VCF) files corresponding to individual wheat chromosomes were downloaded from the Watkins collection repository (https://wwwg2b.com). These chromosome-specific datasets consisted of multi-sample VCF files containing genotypic information for 827 Watkins accessions. All variant positions were reported relative to the *Triticum aestivum* reference genome assembly *IWGSC RefSeq v1.0* (IWGSC 2018).

### Genotype processing and quality control

Chromosome-specific VCF files were converted to PLINK binary format (BED/BIM/FAM) using PLINK v1.9 (Chang et al. 2015). Genotypic data was restricted to 297 individuals present in the phenotypic dataset prior to filtering. Quality control was performed separately for each chromosome. Markers with minor allele frequency lower than 0.02 were removed. SNPs with more than 10% missing genotype calls were excluded, and individuals with more than 10% missing data were removed from the dataset (**Table S2**). Following quality filtering, a two-step linkage disequilibrium (LD) pruning procedure was applied to reduce marker redundancy as described by Cheng et al. (2024) and Wang et al. (2018). First, LD pruning was conducted using a 10 kb sliding window with a step size of one SNP and an r² threshold of 0.8. A second round of pruning was subsequently performed using a 50-SNP window with a step size of one SNP and an r² threshold of 0.8 (**Table S2**). After chromosome-level filtering and pruning, the resulting datasets were merged across all 21 chromosomes to generate a final genome-wide marker dataset for association analysis.

### Population structure and kinship estimation

Population structure was assessed using principal component analysis (PCA) implemented in PLINK v1.9 (Chang et al. 2015). To minimize bias due to local LD, PCA was performed on an LD-pruned subset of markers generated using a 50-SNP sliding window with a step size of five SNPs and an r² threshold of 0.2. The top principal components were extracted from the eigenvector output and incorporated as fixed-effect covariates in subsequent association analyses. Pairwise genetic relatedness among individuals was estimated using the genomic relationship matrix computed in PLINK using the --make-rel square function. The resulting kinship matrix was used in mixed model analyses to account for cryptic relatedness.

### Genome-wide association analysis

Genome-wide association analyses were performed using the PANICLE GWAS framework (https://github.com/jschnable/PANICLE). Genotypic data were supplied in PLINK binary format, and phenotypic data were provided in tab-delimited format with matching individual identifiers. Association analyses were conducted using a general linear model (GLM), a mixed linear model (MLM), and the multi-locus model: Fixed and random model Circulating Probability Unification (FarmCPU) (Liu et al. 2016). Principal components derived from the external PCA were included as covariates to control for population stratification. Internal PCA computation within PANICLE was disabled, as population structure covariates were provided explicitly. Genome-wide significance thresholds were determined using a Bonferroni correction based on the total number of markers in the filtered dataset. Quantile–quantile plots were used to evaluate model performance, and genomic inflation factors were calculated to assess residual population structure. Models with genomic inflation factors approximating unity were considered to provide appropriate control of confounding effects.

### Defining quantitative trait loci (QTL)

Lead SNPs were defined as markers exceeding the Bonferroni-adjusted significance threshold in MLM and/or FarmCPU analyses. To delineate QTL boundaries, local linkage disequilibrium (LD) was calculated for each lead SNP using PLINK within a ±10 Mb physical window. Pairwise r² values were computed between the lead SNP and surrounding markers, and SNPs showing r² ≥0.2 with the lead marker were considered part of the same LD block. The minimum and maximum physical positions of SNPs meeting this LD criterion were used to define the initial QTL interval boundaries for each lead marker. Because multiple significant SNPs frequently occurred within the same genomic region, overlapping LD-defined intervals on the same chromosome were subsequently merged to avoid redundant QTL reporting. Intervals were sorted by chromosomal position and merged when their physical ranges overlapped. The merged interval boundaries were defined by the outermost start and end coordinates across overlapping regions, and all associated lead SNPs were retained as representative markers for the consolidated QTL. This LD-based approach enabled empirical determination of QTL extent while preventing artificial fragmentation of single genomic regions into multiple adjacent QTL. Furthermore, because GWAS was performed independently for six distinct *Pst* isolates using three statistical models, additional criteria were applied to define strain-consistent loci. QTL detected across different strains and/or models were designated as a single locus when they exhibited (i) overlapping LD-defined physical intervals on the same chromosome, (ii) comparable lead SNP positions within the same LD block, and (iii) consistent direction of allelic effect on disease severity. When these criteria were satisfied, signals were consolidated and reported as a single QTL, while retaining strain- and model-specific lead SNP information. Conversely, signals with overlapping physical intervals but distinct LD patterns or inconsistent effect directions were retained as separate QTL.

### *In-silico* physical analysis of mapped QTL

To enable direct comparison with previously reported loci, the physical positions of all identified QTL were converted to the latest wheat reference assembly, *IWGSC RefSeq v2.1*. Genomic coordinates were harmonized to ensure consistency across analyses and downstream comparisons. The resulting QTL intervals were then compared based on their physical positions with all formally designated and cloned *Yr* genes, previously reported stripe rust QTL, and QTL-rich genomic clusters summarized in Tong et al. (2024). Co-localization was assessed by examining overlap between the LD-defined QTL intervals and the published physical coordinates of known loci on the same chromosome.

### Candidate gene prediction

Candidate genes within LD-defined QTL intervals were identified/predicted using the *Triticum aestivum IWGSC RefSeq v1.0* gene annotation obtained from Ensembl Plants release 62. Gene features were extracted from the GFF3 annotation file and converted to BED format. LD-defined QTL intervals were intersected with gene coordinates using BEDTools (Quinlan and Hall 2010). Genes whose genomic coordinates overlapped QTL intervals were considered positional candidate genes and were retained for further functional annotation and biological interpretation.

### Haplotype analysis

To further characterize allelic variation within identified QTL, haplotype analysis was conducted for all the high confidence resistance loci. For each QTL, SNPs within the LD-defined interval surrounding the lead SNP were extracted from the genotype dataset using PLINK v1.9. Pairwise linkage disequilibrium (LD) between the lead SNP and surrounding markers was calculated using the --r2 function in PLINK within a ±30 Mb window. SNPs showing strong LD with the lead marker were retained as candidate markers for haplotype construction. Genotype dosages for the selected SNPs were extracted using the --recode A option in PLINK, and haplotype patterns for each accession were generated by concatenating allele dosages across the selected SNPs. Haplotypes containing missing or heterozygous genotype calls were excluded to retain only homozygous haplotype configurations. Haplotype allele states were subsequently converted to nucleotide calls (A/T/G/C) based on allele information from the corresponding BIM file. For each haplotype, the number of accessions carrying the haplotype, mean infection score, and standard deviation were calculated. Accessions sharing the same haplotype were grouped to evaluate phenotypic differences among haplotypes. Statistical significance of phenotypic differences between haplotypes was assessed using the non-parametric Mann–Whitney U test, with a significance threshold of *P* <0.05.

## Results

### Phenotypic variation of Watkins landraces to six *Pst* isolates

Stripe rust response of the 297 Watkins landraces was evaluated against six *Pst* isolates (W034, W001, T022, W056, W031, and W057) representing three diverse lineages that revealed substantial phenotypic variation among genotypes and isolates (**Fig. 1; Table S1**). Some landraces maintained consistently IT across isolates, whereas others exhibited contrasting reactions depending on the isolate tested. WATDE0042, WATDE0671, WATDE0694, and WATDE0776 maintained IT <=2 for each isolate (**Fig. 1a; Table S1**). In contrast, several genotypes showed isolate-dependent responses. For example, WATDE0031, WATDE0080, WATDE0081, WATDE0919, and WATDE0392 exhibited high IT of >=7) to W034 and low severity to all the other isolates. These contrasting patterns indicated that genotype performance was not uniform across isolates and that screening with multiple *Pst* isolates captured additional phenotypic differentiation within the panel.

**Figure 1.**
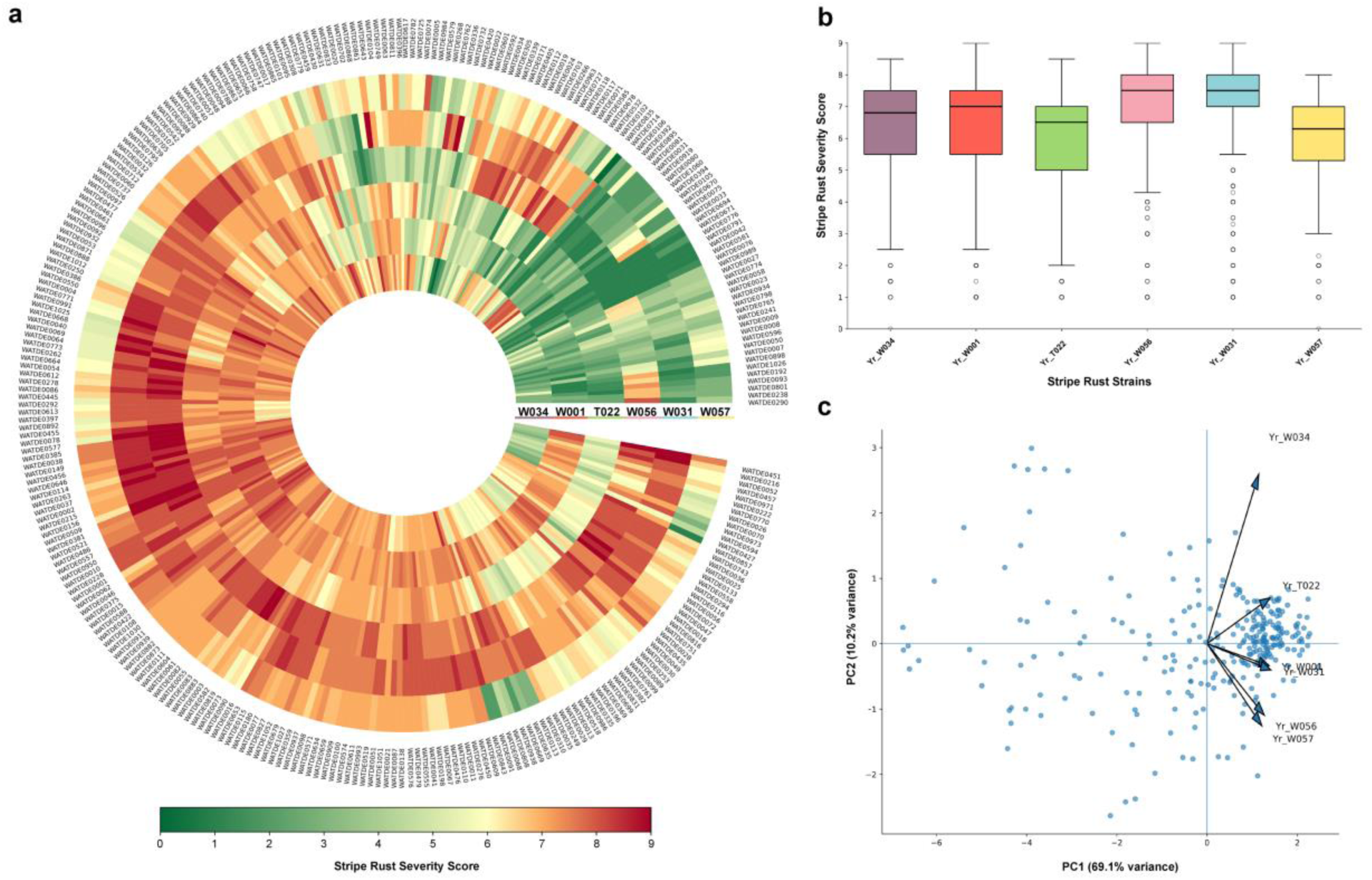
Phenotypic variation and multivariate structure of stripe rust responses in the Watkins landrace panel across six *Puccinia striiformis* f. sp. *tritici* isolates. (a) Circular heatmap of disease severity scores (0–9) for 297 Watkins landraces evaluated against isolates W034, W001, T022, W056, W031 and W057. Each ring represents one isolate and each segment an accession; colors range from green (low severity) to red (high severity). (b) Boxplots showing severity score distributions per isolate; boxes indicate interquartile range, horizontal lines medians, whiskers ranges (excluding outliers), and points outliers. (c) Principal component analysis of phenotypic responses. PC1 (69.1%) reflects overall disease severity, and PC2 (10.2%) captures isolate-specific variation. Arrows denote isolate loadings and points represent individual accessions.

IT distributions were continuous for all isolates, without clear separation into discrete resistant and susceptible classes (**Fig. 1b**). Median IT differed among isolates, with W056 and W031 showing comparatively higher central tendencies, whereas T022 and W034 displayed broader dispersion of genotype responses.

Principal component analysis on phenotypic data further resolved the structure of phenotypic variation (**Fig. 1c**). PC1 explained 69.1% of the total phenotypic variance and separated genotypes along a general severity gradient, while PC2 accounted for 10.2% of the variance and distinguished isolates based on differences in response patterns. Loading vectors showed close alignment between W056 and W057, as well as between W001 and W031, whereas W034 was separated along PC2 relative to the other isolates. The clustering of genotypes along PC1 suggests that a major portion of phenotypic variation represents overall disease severity across isolates, with additional components differentiating isolate-specific responses.

### Genotypic dataset composition and filtering

Whole-genome SNP data was processed on a chromosome-by-chromosome basis. After filtering for minor allele frequency (MAF ≥0.02), marker missingness (≤10%), and individual missingness (≤10%), a total of 73,665,006 high-quality SNPs were retained across all 21 wheat chromosomes (**Table S2**). The final dataset included 297 accessions, with no individuals removed after sample-level filtering. Markers were distributed across the A, B, and D sub genomes, with the B genome contributing the largest proportion of polymorphic loci and the D genome showing comparatively lower diversity (**Table S2**).

To reduce marker redundancy for downstream analyses, a two-step LD pruning procedure was implemented. Initial pruning using a 10 kb sliding window (r² ≤0.8) was followed by a second pruning step using a 50-SNP window (r² ≤0.8), yielding 5,652,314 SNPs (**Table S2**). This strategy substantially reduced local marker redundancy while retaining genome-wide coverage. The resulting pruned dataset was used for population structure analysis.

### Population structure and genomic relatedness of the Watkins accessions

Principal component analysis based on the LD-pruned marker set revealed moderate genetic structure within the Watkins panel (**Fig. 2**). PC1 explained 13.6% of the total genetic variance, while PC2 and PC3 accounted for 8.6% and 7.5%, respectively. Together, the first three components captured 29.7% of the total genomic variation.

**Figure 2.**
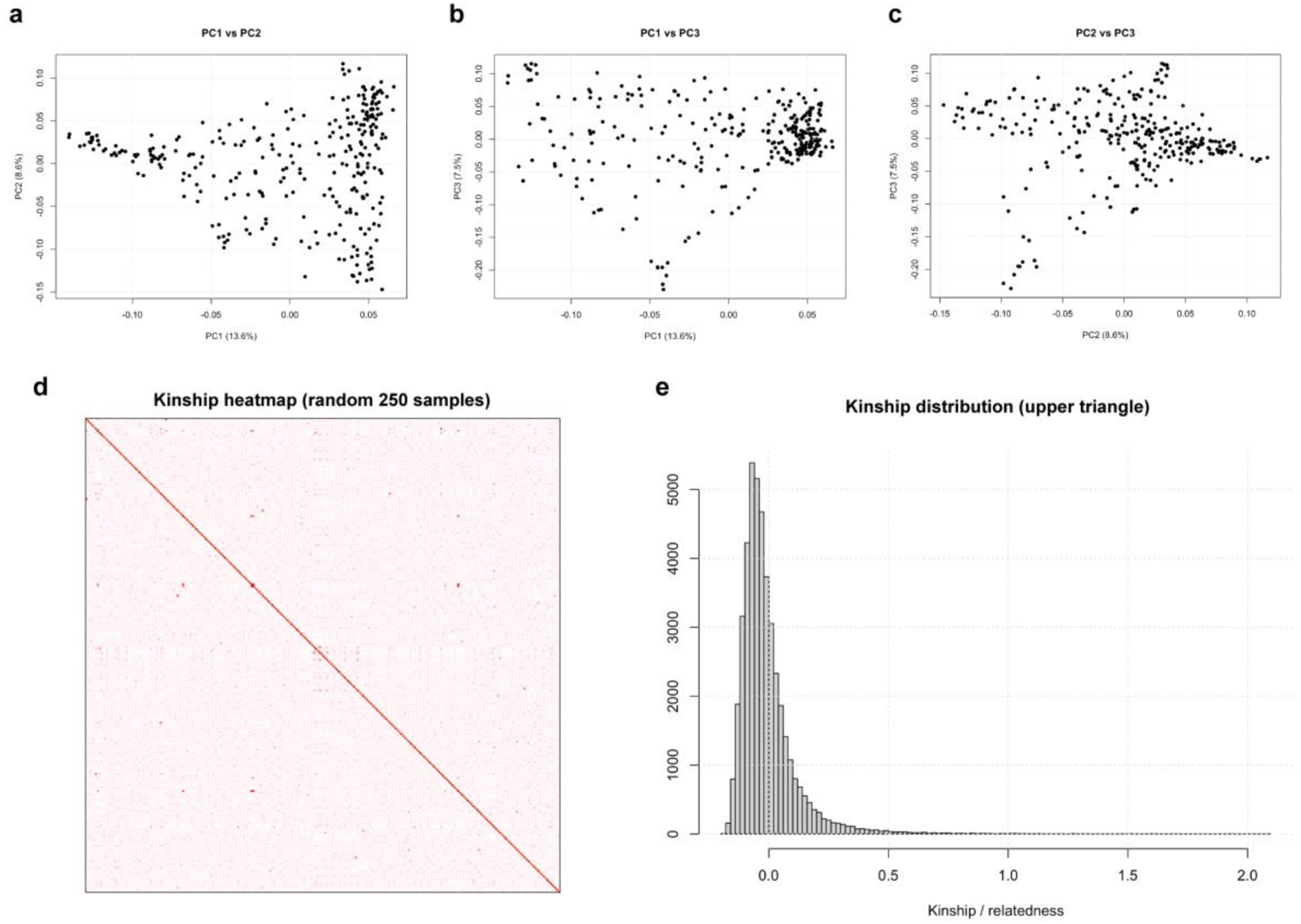
Population structure and kinship relationships in the Watkins wheat landrace panel based on LD-pruned SNP markers. (a) Principal component analysis (PCA) showing the distribution of accessions along PC1 (13.6% variance explained) and PC2 (8.6%). (b) PCA plot of PC1 (13.6%) versus PC3 (7.5%), illustrating additional axes of genetic variation. (c) PCA plot of PC2 (8.6%) versus PC3 (7.5%), highlighting subtle genetic differentiation among accessions. (d) Kinship heatmap based on the realized genomic relationship matrix (random subset of 250 accessions), with color intensity reflecting pairwise relatedness. (e) Distribution of pairwise kinship coefficients (upper triangle of the matrix), indicating that most accession pairs exhibit low relatedness with a right-skewed distribution toward higher values.

Visualization of PC1 versus PC2 indicated continuous genetic differentiation rather than discrete, well-separated clusters (**Fig. 2a**). Accessions were broadly distributed along PC1, with values ranging approximately from −0.14 to 0.06, while PC2 values ranged from approximately −0.15 to 0.12. Although partial grouping was evident along PC1, particularly toward positive PC1 values, clear-cut subpopulation boundaries were not observed. Similar patterns were evident in PC1 versus PC3 and PC2 versus PC3 projections (**Fig. 2b, c**), where gradual gradients of variation were observed instead of sharply defined clusters.

Estimation of pairwise genomic relationships revealed generally low to moderate relatedness among accessions (**Fig. 2 d, e**). The majority of pairwise kinship coefficients were centered near zero, with a small number of higher values reflecting shared ancestry among closely related accessions. The absence of widespread high kinship values indicates limited redundancy within the panel and supports its suitability for genome-wide association analysis.

### Quantitative trait nucleotides (QTNs)

Genome-wide association analyses were performed for six *Pst* isolates using three statistical models: GLM, MLM, and FarmCPU. Quantile–quantile plots consistently revealed substantial genomic inflation under the GLM model (λ >1.3 across strains), indicating insufficient control of population structure and relatedness. Consequently, GLM-derived associations were not considered reliable and were excluded from further interpretation. All subsequent reporting of significant loci was therefore based exclusively on the MLM and FarmCPU models.

Using a Bonferroni-adjusted significance threshold (α = 0.05/5,657,375 markers), a total of 1,602 significant QTNs were identified across the six stripe rust strains when combining results from MLM and FarmCPU (**Table S3**). The number of QTNs detected varied markedly among isolates. W056 exhibited the highest number of associations, with 1,118 QTNs identified by MLM and 9 by FarmCPU (1,127 total) (**Fig. 3; Table S3**). W031 showed 250 QTNs detected by MLM and 10 by FarmCPU (260 total) (**Fig. S1; Table S3**). W057 yielded 123 QTNs from MLM and 5 from FarmCPU (128 total) (**Fig. S2; Table S3**), whereas W034 displayed 64 QTNs identified by MLM and 5 by FarmCPU (69 total) (**Fig. S3; Table S3**). W001 showed 6 QTNs from MLM and 8 from FarmCPU (14 total) (**Fig. S4; Table S3**), while T022 exhibited only 4 QTNs (**Fig S5; Table S3**), all detected exclusively by FarmCPU.

**Figure 3.**
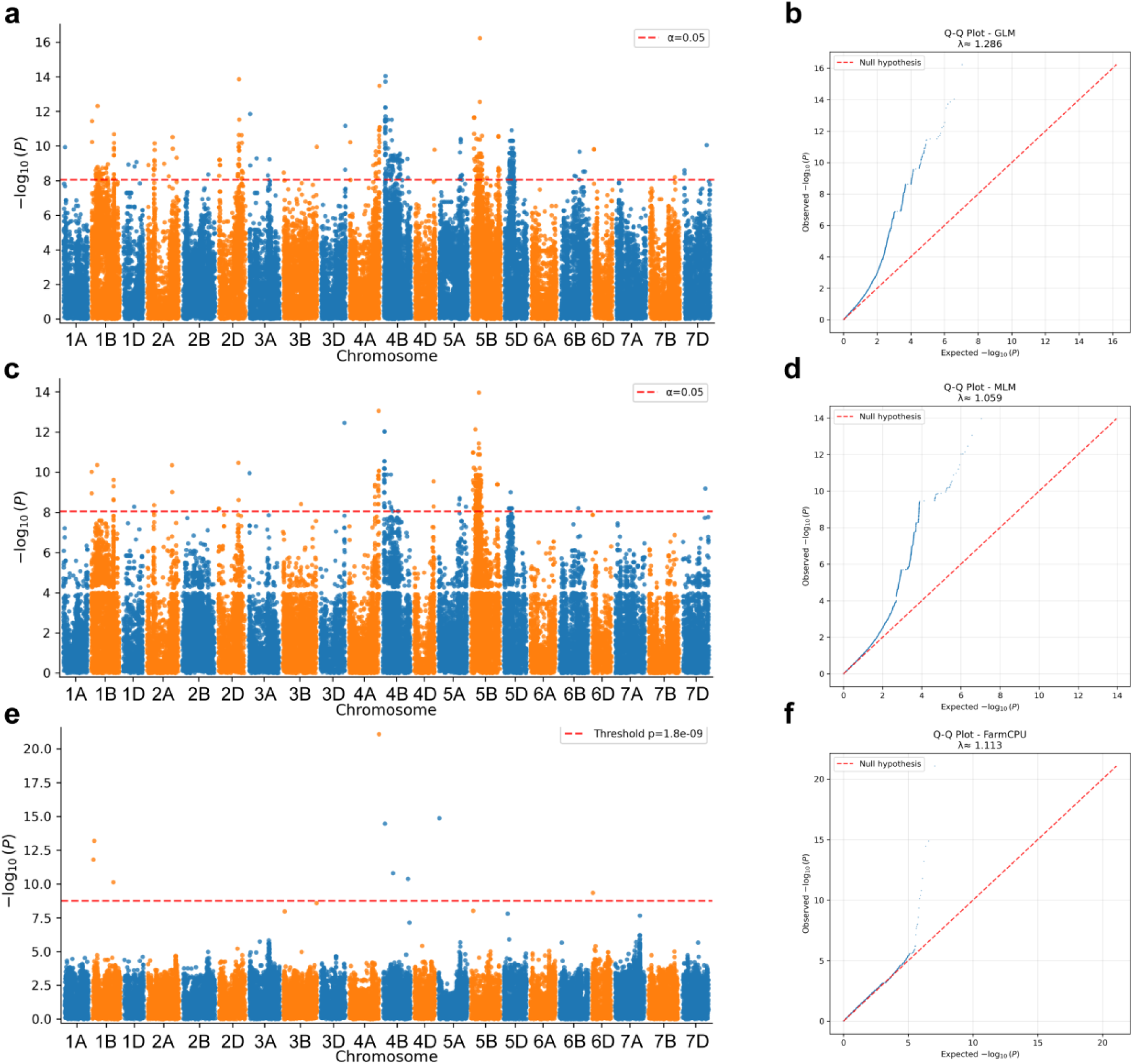
Genome-wide association analysis for stripe rust isolate W056 using GLM, MLM, and FarmCPU models. (a) Manhattan plot from the GLM analysis showing genome-wide marker–trait associations across the 21 wheat chromosomes. The red dashed line indicates the significance threshold (α = 0.05). (b) Q–Q plot for the GLM model illustrating the deviation of observed from expected –log₁₀(P) values, with an inflation factor (λ) of 1.286. (c) Manhattan plot from the MLM analysis incorporating population structure and kinship. The red dashed line denotes the significance threshold (α = 0.05). (d) Q–Q plot for the MLM model (λ = 1.059), indicating improved control of false positives relative to GLM. (e) Manhattan plot from the FarmCPU analysis, with the genome-wide significance threshold (P = 1.8 × 10⁻⁹) shown by the red dashed line. (f) Q–Q plot for the FarmCPU model (λ = 1.113), demonstrating effective control of population structure and cryptic relatedness while retaining significant associations.

**Figure 4.**
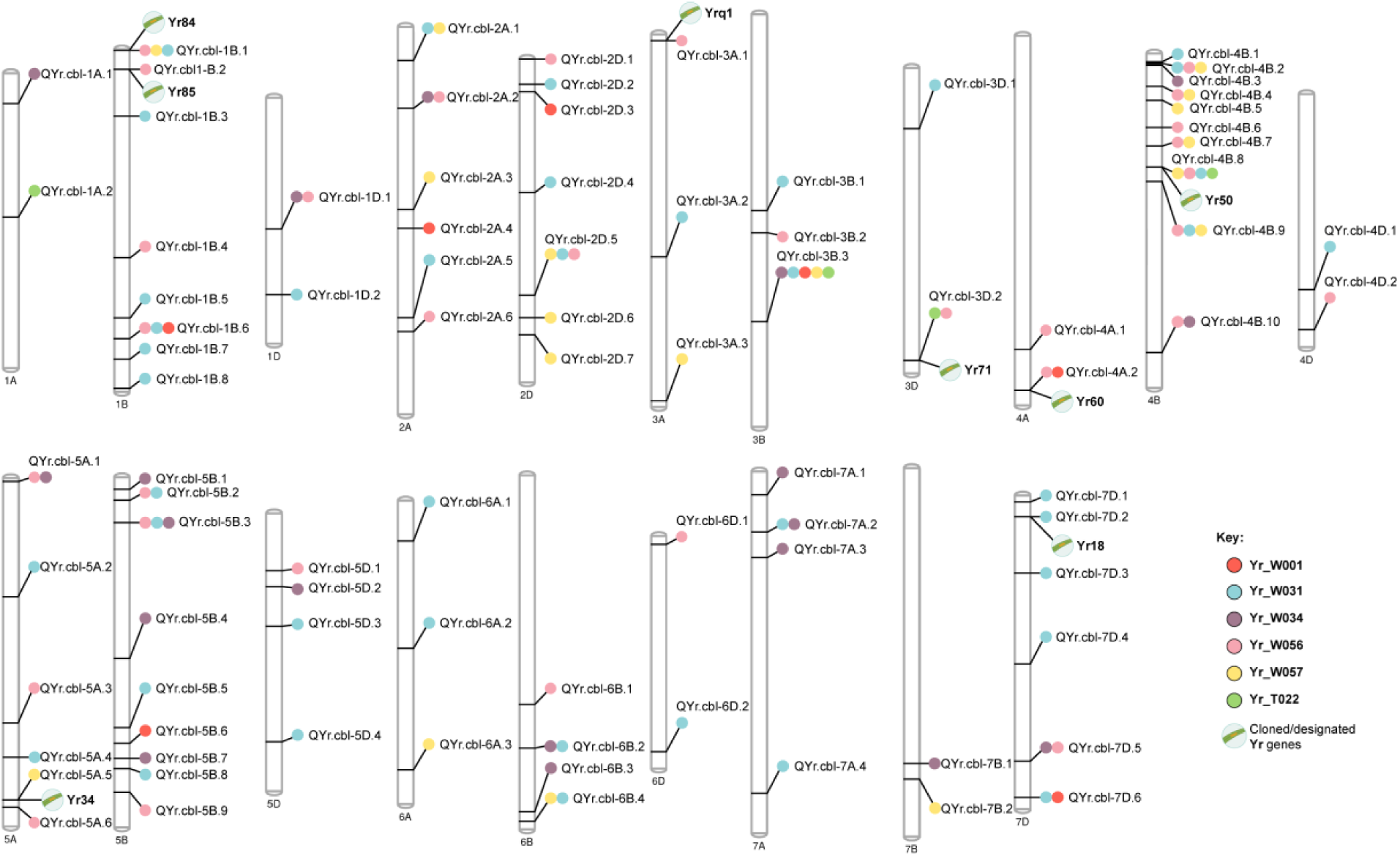
Chromosomal distribution of stripe rust resistance QTL identified in the Watkins panel and their co-localization with previously reported *Yr* genes.

### QTL identification

Using the LD-based interval delineation and consolidation strategy described in the ‘Materials and methods’ section, a total of 87 non-redundant QTL were identified for stripe rust response across six *Pst* isolates (**Table S4; Table 2**). Physical intervals of the QTL varied considerably in size, reflecting local linkage disequilibrium structure and marker density.

**Table 2.**
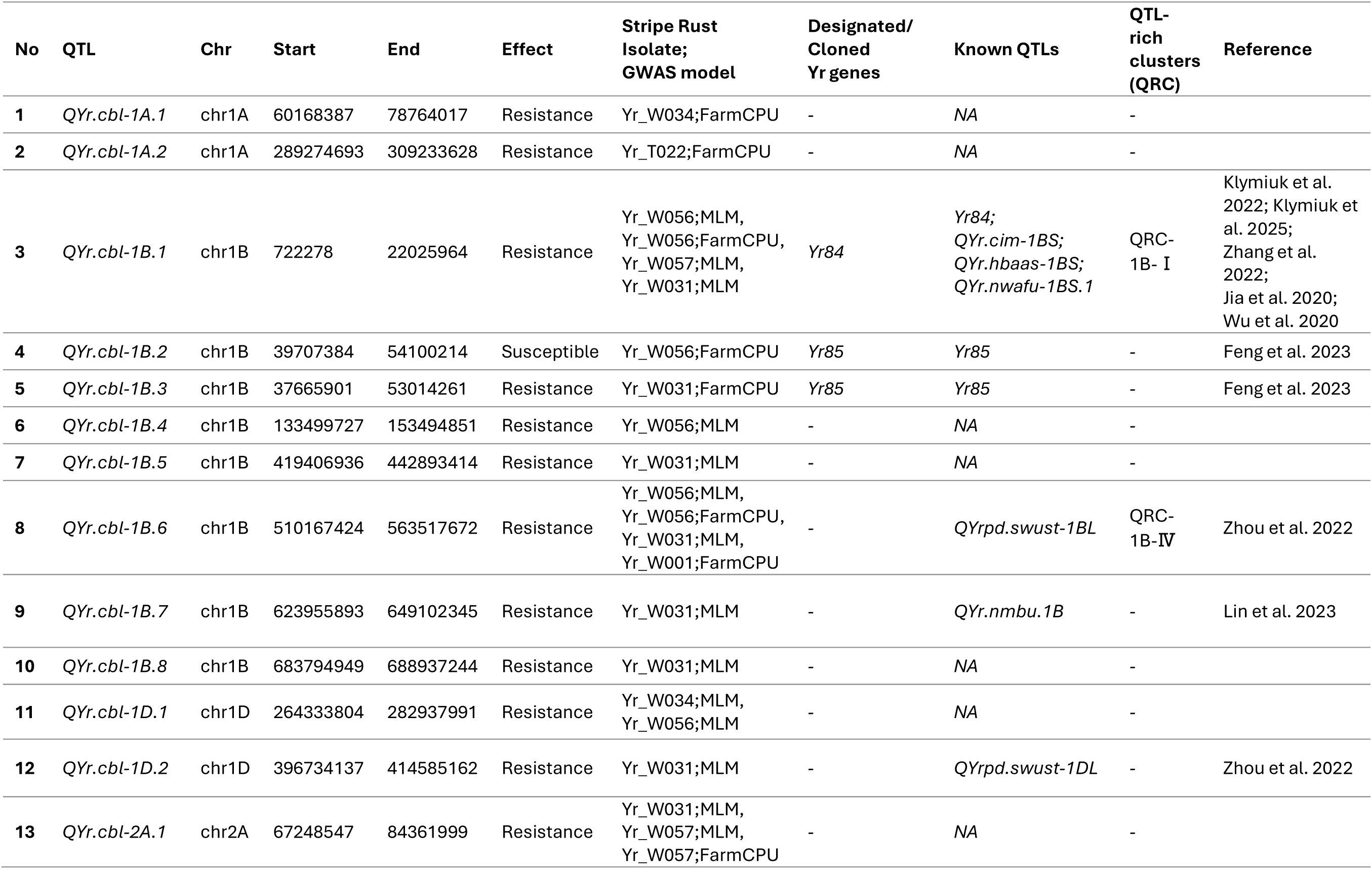
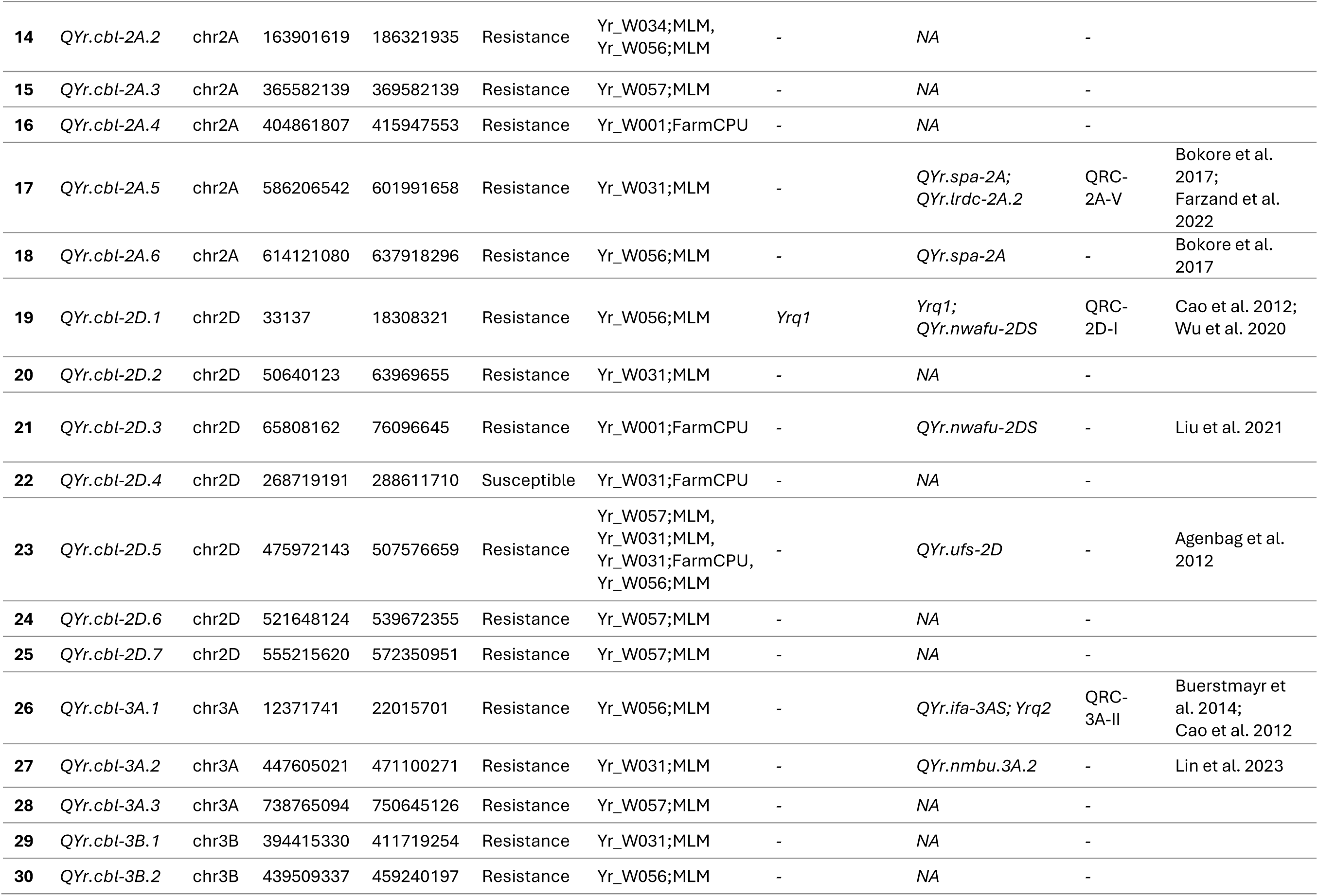
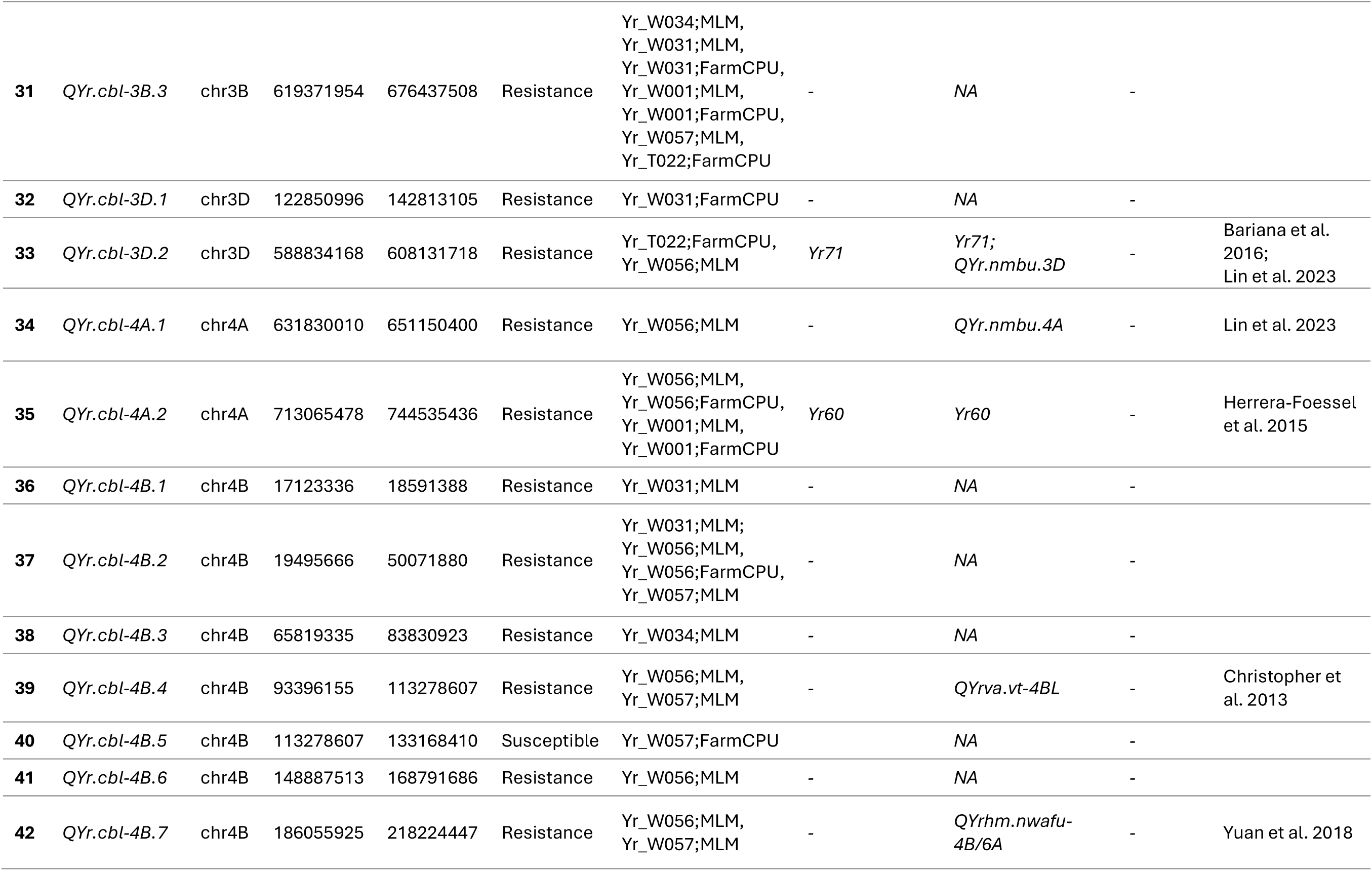
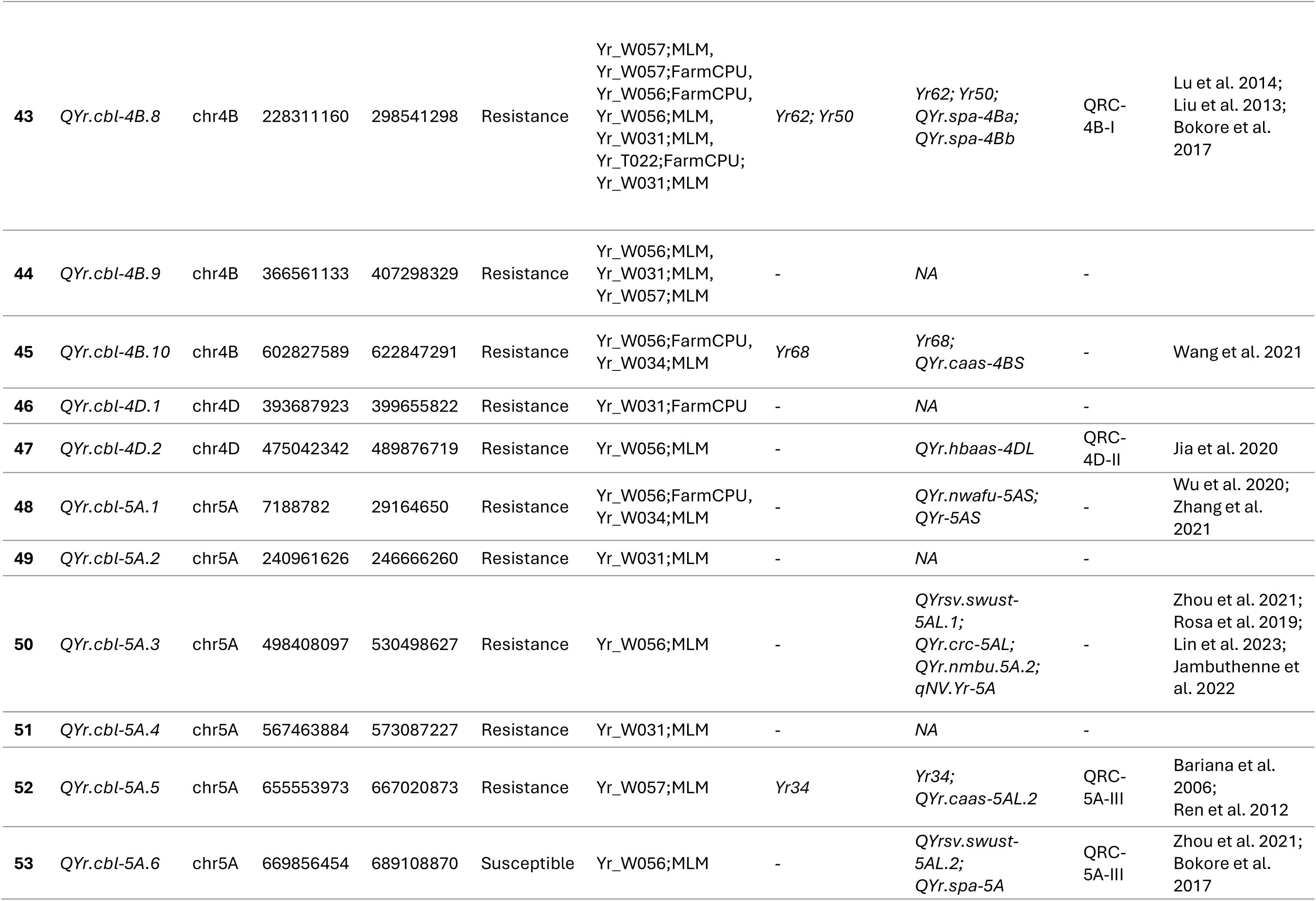
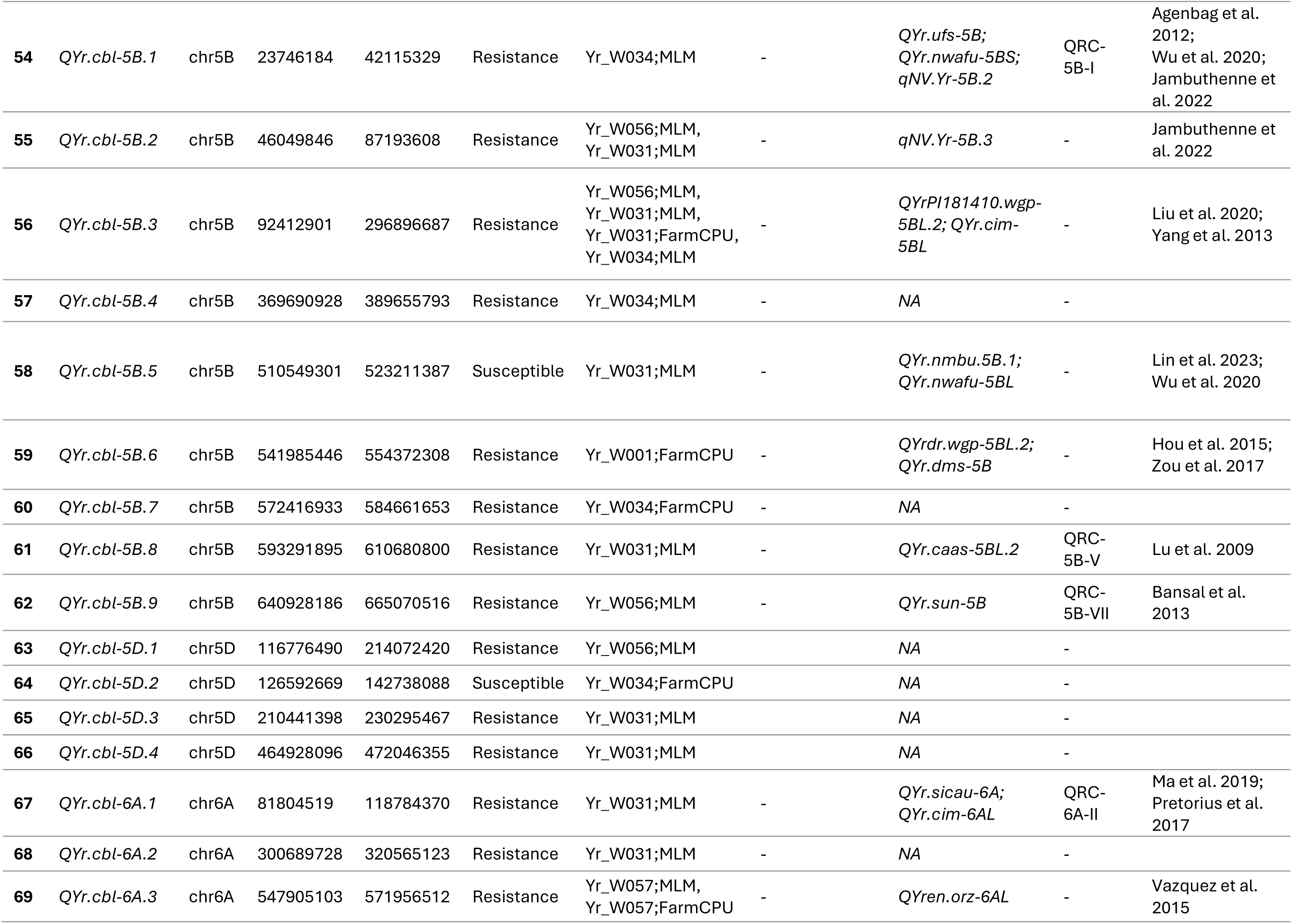
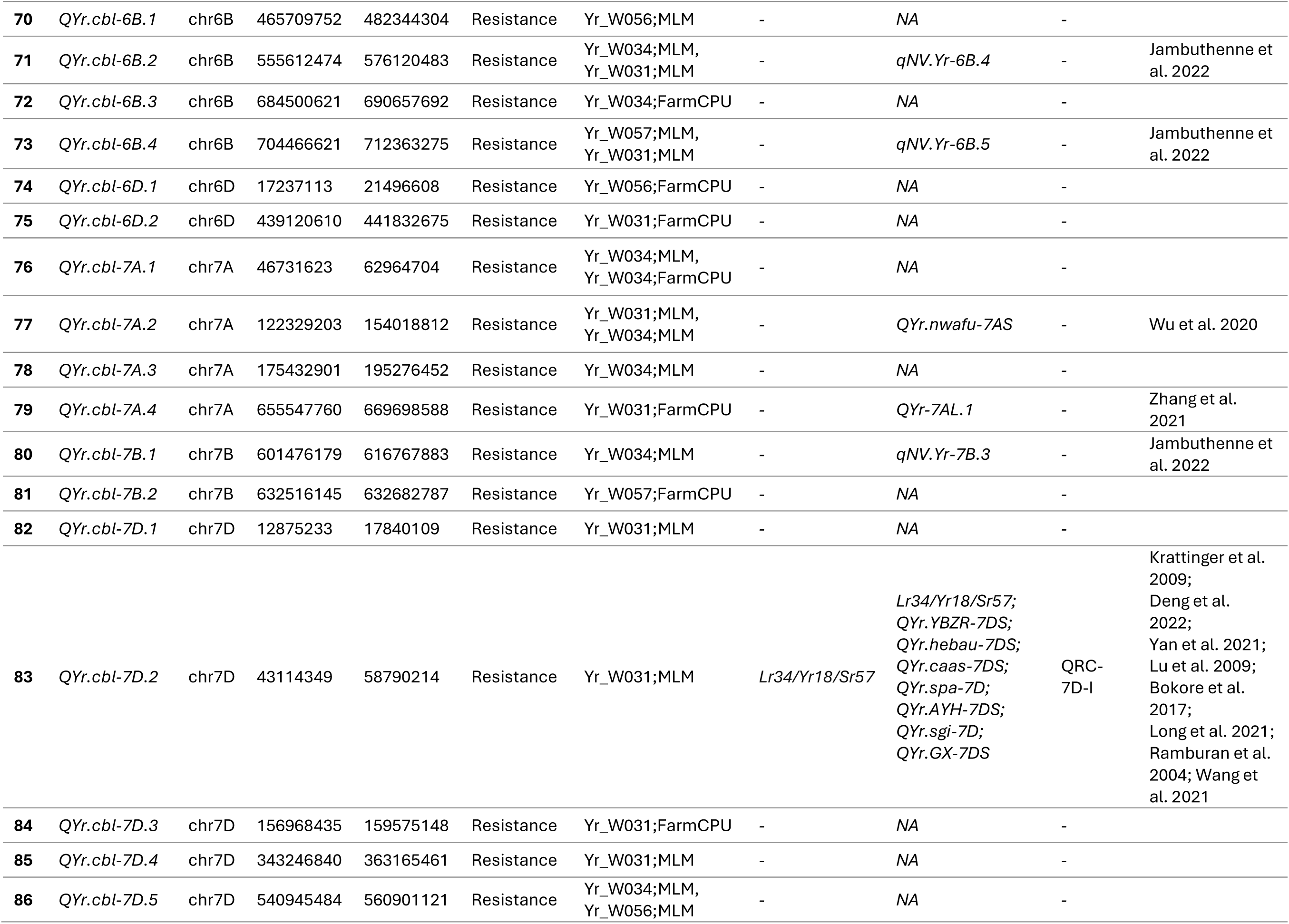
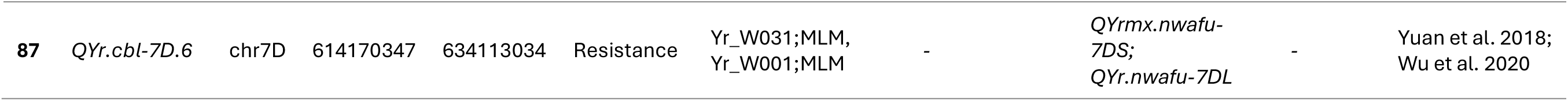
Genome-wide stripe rust QTL detected across six diverse isolates and their co-localization with known *Yr* genes and QTL.

| No | QTL | Chr | Start | End | Effect | Stripe Rust Isolate;<br>GWAS model | Designated/<br>Cloned<br>Yr genes | Known QTLs | QTL-rich<br>clusters<br>(QRC) | Reference |
| --- | --- | --- | --- | --- | --- | --- | --- | --- | --- | --- |
| 1 | *QYr.cbl-1A.1* | chr1A | 60168387 | 78764017 | Resistance | Yr_W034;FarmCPU | - | NA | - |  |
| 2 | *QYr.cbl-1A.2* | chr1A | 289274693 | 309233628 | Resistance | Yr_T022;FarmCPU | - | NA | - |  |
| 3 | *QYr.cbl-1B.1* | chr1B | 722278 | 22025964 | Resistance | Yr_W056;MLM,<br>Yr_W056;FarmCPU,<br>Yr_W057;MLM,<br>Yr_W031;MLM | *Yr84* | *Yr84*;<br>*QYr.cim-1BS*;<br>*QYr.hbaas-1BS*;<br>*QYr.nwafu-1BS.1* | QRC-1B- I | Klymiuk et al.<br>2022; Klymiuk et<br>al. 2025;<br>Zhang et al.<br>2022;<br>Jia et al. 2020;<br>Wu et al. 2020 |
| 4 | *QYr.cbl-1B.2* | chr1B | 39707384 | 54100214 | Susceptible | Yr_W056;FarmCPU | *Yr85* | *Yr85* | - | Feng et al. 2023 |
| 5 | *QYr.cbl-1B.3* | chr1B | 37665901 | 53014261 | Resistance | Yr_W031;FarmCPU | *Yr85* | *Yr85* | - | Feng et al. 2023 |
| 6 | *QYr.cbl-1B.4* | chr1B | 133499727 | 153494851 | Resistance | Yr_W056;MLM | - | NA | - |  |
| 7 | *QYr.cbl-1B.5* | chr1B | 419406936 | 442893414 | Resistance | Yr_W031;MLM | - | NA | - |  |
| 8 | *QYr.cbl-1B.6* | chr1B | 510167424 | 563517672 | Resistance | Yr_W056;MLM,<br>Yr_W056;FarmCPU,<br>Yr_W031;MLM,<br>Yr_W001;FarmCPU | - | *QYrpd.swust-1BL* | QRC-1B-IV | Zhou et al. 2022 |
| 9 | *QYr.cbl-1B.7* | chr1B | 623955893 | 649102345 | Resistance | Yr_W031;MLM | - | *QYr.nmbu.1B* | - | Lin et al. 2023 |
| 10 | *QYr.cbl-1B.8* | chr1B | 683794949 | 688937244 | Resistance | Yr_W031;MLM | - | NA | - |  |
| 11 | *QYr.cbl-1D.1* | chr1D | 264333804 | 282937991 | Resistance | Yr_W034;MLM,<br>Yr_W056;MLM | - | NA | - |  |
| 12 | *QYr.cbl-1D.2* | chr1D | 396734137 | 414585162 | Resistance | Yr_W031;MLM | - | *QYrpd.swust-1DL* | - | Zhou et al. 2022 |
| 13 | *QYr.cbl-2A.1* | chr2A | 67248547 | 84361999 | Resistance | Yr_W031;MLM,<br>Yr_W057;MLM,<br>Yr_W057;FarmCPU | - | NA | - |  |
| 14 | *QYr.cbl-2A.2* | chr2A | 163901619 | 186321935 | Resistance | Yr_W034;MLM,<br>Yr_W056;MLM | - | NA | - |  |
| 15 | *QYr.cbl-2A.3* | chr2A | 365582139 | 369582139 | Resistance | Yr_W057;MLM | - | NA | - |  |
| 16 | *QYr.cbl-2A.4* | chr2A | 404861807 | 415947553 | Resistance | Yr_W001;FarmCPU | - | NA | - |  |
| 17 | *QYr.cbl-2A.5* | chr2A | 586206542 | 601991658 | Resistance | Yr_W031;MLM | - | *QYr.spa-2A;*<br>*QYr.lrdc-2A.2* | QRC-2A-V | Bokore et al.<br>2017;<br>Farzand et al.<br>2022 |
| 18 | *QYr.cbl-2A.6* | chr2A | 614121080 | 637918296 | Resistance | Yr_W056;MLM | - | *QYr.spa-2A* | - | Bokore et al.<br>2017 |
| 19 | *QYr.cbl-2D.1* | chr2D | 33137 | 18308321 | Resistance | Yr_W056;MLM | *Yrq1* | *Yrq1;*<br>*QYr.nwafu-2DS* | QRC-2D-I | Cao et al. 2012;<br>Wu et al. 2020 |
| 20 | *QYr.cbl-2D.2* | chr2D | 50640123 | 63969655 | Resistance | Yr_W031;MLM | - | NA | - |  |
| 21 | *QYr.cbl-2D.3* | chr2D | 65808162 | 76096645 | Resistance | Yr_W001;FarmCPU | - | *QYr.nwafu-2DS* | - | Liu et al. 2021 |
| 22 | *QYr.cbl-2D.4* | chr2D | 268719191 | 288611710 | Susceptible | Yr_W031;FarmCPU | - | NA | - |  |
| 23 | *QYr.cbl-2D.5* | chr2D | 475972143 | 507576659 | Resistance | Yr_W057;MLM,<br>Yr_W031;MLM,<br>Yr_W031;FarmCPU,<br>Yr_W056;MLM | - | *QYr.ufs-2D* | - | Agenbag et al.<br>2012 |
| 24 | *QYr.cbl-2D.6* | chr2D | 521648124 | 539672355 | Resistance | Yr_W057;MLM | - | NA | - |  |
| 25 | *QYr.cbl-2D.7* | chr2D | 555215620 | 572350951 | Resistance | Yr_W057;MLM | - | NA | - |  |
| 26 | *QYr.cbl-3A.1* | chr3A | 12371741 | 22015701 | Resistance | Yr_W056;MLM | - | *QYr.ifa-3AS; Yrq2* | QRC-3A-II | Buerstmayr et<br>al. 2014;<br>Cao et al. 2012 |
| 27 | *QYr.cbl-3A.2* | chr3A | 447605021 | 471100271 | Resistance | Yr_W031;MLM | - | *QYr.nmbu.3A.2* | - | Lin et al. 2023 |
| 28 | *QYr.cbl-3A.3* | chr3A | 738765094 | 750645126 | Resistance | Yr_W057;MLM | - | NA | - |  |
| 29 | *QYr.cbl-3B.1* | chr3B | 394415330 | 411719254 | Resistance | Yr_W031;MLM | - | NA | - |  |
| 30 | *QYr.cbl-3B.2* | chr3B | 439509337 | 459240197 | Resistance | Yr_W056;MLM | - | NA | - |  |
| 31 | *QYr.cbl-3B.3* | chr3B | 619371954 | 676437508 | Resistance | Yr_W034;MLM,<br>Yr_W031;MLM,<br>Yr_W031;FarmCPU,<br>Yr_W001;MLM,<br>Yr_W001;FarmCPU,<br>Yr_W057;MLM,<br>Yr_T022;FarmCPU | - | NA | - |  |
| 32 | *QYr.cbl-3D.1* | chr3D | 122850996 | 142813105 | Resistance | Yr_W031;FarmCPU | - | NA | - |  |
| 33 | *QYr.cbl-3D.2* | chr3D | 588834168 | 608131718 | Resistance | Yr_T022;FarmCPU,<br>Yr_W056;MLM | *Yr71* | *Yr71;*<br>*QYr.nmbu.3D* | - | Bariana et al.<br>2016;<br>Lin et al. 2023 |
| 34 | *QYr.cbl-4A.1* | chr4A | 631830010 | 651150400 | Resistance | Yr_W056;MLM | - | *QYr.nmbu.4A* | - | Lin et al. 2023 |
| 35 | *QYr.cbl-4A.2* | chr4A | 713065478 | 744535436 | Resistance | Yr_W056;MLM,<br>Yr_W056;FarmCPU,<br>Yr_W001;MLM,<br>Yr_W001;FarmCPU | *Yr60* | *Yr60* | - | Herrera-Foessel<br>et al. 2015 |
| 36 | *QYr.cbl-4B.1* | chr4B | 17123336 | 18591388 | Resistance | Yr_W031;MLM | - | NA | - |  |
| 37 | *QYr.cbl-4B.2* | chr4B | 19495666 | 50071880 | Resistance | Yr_W031;MLM;<br>Yr_W056;MLM,<br>Yr_W056;FarmCPU,<br>Yr_W057;MLM | - | NA | - |  |
| 38 | *QYr.cbl-4B.3* | chr4B | 65819335 | 83830923 | Resistance | Yr_W034;MLM | - | NA | - |  |
| 39 | *QYr.cbl-4B.4* | chr4B | 93396155 | 113278607 | Resistance | Yr_W056;MLM,<br>Yr_W057;MLM | - | *QYrva.vt-4BL* | - | Christopher et<br>al. 2013 |
| 40 | *QYr.cbl-4B.5* | chr4B | 113278607 | 133168410 | Susceptible | Yr_W057;FarmCPU |  | NA | - |  |
| 41 | *QYr.cbl-4B.6* | chr4B | 148887513 | 168791686 | Resistance | Yr_W056;MLM | - | NA | - |  |
| 42 | *QYr.cbl-4B.7* | chr4B | 186055925 | 218224447 | Resistance | Yr_W056;MLM,<br>Yr_W057;MLM | - | *QYrhm.nwafu-*<br>*4B/6A* | - | Yuan et al. 2018 |
| 43 | *QYr.cbl-4B.8* | chr4B | 228311160 | 298541298 | Resistance | Yr_W057;MLM,<br>Yr_W057;FarmCPU,<br>Yr_W056;FarmCPU,<br>Yr_W056;MLM,<br>Yr_W031;MLM,<br>Yr_T022;FarmCPU;<br>Yr_W031;MLM | Yr62; Yr50 | Yr62; Yr50;<br>*QYr.spa-4Ba*;<br>*QYr.spa-4Bb* | QRC-4B-I | Lu et al. 2014;<br>Liu et al. 2013;<br>Bokore et al.<br>2017 |
| 44 | *QYr.cbl-4B.9* | chr4B | 366561133 | 407298329 | Resistance | Yr_W056;MLM,<br>Yr_W031;MLM,<br>Yr_W057;MLM | - | NA | - |  |
| 45 | *QYr.cbl-4B.10* | chr4B | 602827589 | 622847291 | Resistance | Yr_W056;FarmCPU,<br>Yr_W034;MLM | Yr68 | Yr68;<br>*QYr.caas-4BS* | - | Wang et al. 2021 |
| 46 | *QYr.cbl-4D.1* | chr4D | 393687923 | 399655822 | Resistance | Yr_W031;FarmCPU | - | NA | - |  |
| 47 | *QYr.cbl-4D.2* | chr4D | 475042342 | 489876719 | Resistance | Yr_W056;MLM | - | *QYr.hbaas-4DL* | QRC-4D-II | Jia et al. 2020 |
| 48 | *QYr.cbl-5A.1* | chr5A | 7188782 | 29164650 | Resistance | Yr_W056;FarmCPU,<br>Yr_W034;MLM | - | *QYr.nwafu-5AS*;<br>*QYr-5AS* | - | Wu et al. 2020;<br>Zhang et al.<br>2021 |
| 49 | *QYr.cbl-5A.2* | chr5A | 240961626 | 246666260 | Resistance | Yr_W031;MLM | - | NA | - |  |
| 50 | *QYr.cbl-5A.3* | chr5A | 498408097 | 530498627 | Resistance | Yr_W056;MLM | - | *QYrsv.swust-5AL.1*;<br>*QYr.crc-5AL*;<br>*QYr.nmbu.5A.2*;<br>*qNV.Yr-5A* | - | Zhou et al. 2021;<br>Rosa et al. 2019;<br>Lin et al. 2023;<br>Jambuthenne et<br>al. 2022 |
| 51 | *QYr.cbl-5A.4* | chr5A | 567463884 | 573087227 | Resistance | Yr_W031;MLM | - | NA | - |  |
| 52 | *QYr.cbl-5A.5* | chr5A | 655553973 | 667020873 | Resistance | Yr_W057;MLM | Yr34 | Yr34;<br>*QYr.caas-5AL.2* | QRC-5A-III | Bariana et al.<br>2006;<br>Ren et al. 2012 |
| 53 | *QYr.cbl-5A.6* | chr5A | 669856454 | 689108870 | Susceptible | Yr_W056;MLM | - | *QYrsv.swust-5AL.2*;<br>*QYr.spa-5A* | QRC-5A-III | Zhou et al. 2021;<br>Bokore et al.<br>2017 |
| 54 | *QYr.cbl-5B.1* | chr5B | 23746184 | 42115329 | Resistance | Yr_W034;MLM | - | *QYr.ufs-5B;*<br>*QYr.nwafu-5BS;*<br>*qNV.Yr-5B.2* | QRC-5B-I | Agenbag et al. 2012;<br>Wu et al. 2020;<br>Jambuthenne et al. 2022 |
| 55 | *QYr.cbl-5B.2* | chr5B | 46049846 | 87193608 | Resistance | Yr_W056;MLM,<br>Yr_W031;MLM | - | *qNV.Yr-5B.3* | - | Jambuthenne et al. 2022 |
| 56 | *QYr.cbl-5B.3* | chr5B | 92412901 | 296896687 | Resistance | Yr_W056;MLM,<br>Yr_W031;MLM,<br>Yr_W031;FarmCPU,<br>Yr_W034;MLM | - | *QYrPI181410.wgp-5BL.2;*<br>*QYr.cim-5BL* | - | Liu et al. 2020;<br>Yang et al. 2013 |
| 57 | *QYr.cbl-5B.4* | chr5B | 369690928 | 389655793 | Resistance | Yr_W034;MLM | - | NA | - |  |
| 58 | *QYr.cbl-5B.5* | chr5B | 510549301 | 523211387 | Susceptible | Yr_W031;MLM | - | *QYr.nmbu.5B.1;*<br>*QYr.nwafu-5BL* | - | Lin et al. 2023;<br>Wu et al. 2020 |
| 59 | *QYr.cbl-5B.6* | chr5B | 541985446 | 554372308 | Resistance | Yr_W001;FarmCPU | - | *QYrdr.wgp-5BL.2;*<br>*QYr.dms-5B* | - | Hou et al. 2015;<br>Zou et al. 2017 |
| 60 | *QYr.cbl-5B.7* | chr5B | 572416933 | 584661653 | Resistance | Yr_W034;FarmCPU | - | NA | - |  |
| 61 | *QYr.cbl-5B.8* | chr5B | 593291895 | 610680800 | Resistance | Yr_W031;MLM | - | *QYr.caas-5BL.2* | QRC-5B-V | Lu et al. 2009 |
| 62 | *QYr.cbl-5B.9* | chr5B | 640928186 | 665070516 | Resistance | Yr_W056;MLM | - | *QYr.sun-5B* | QRC-5B-VII | Bansal et al. 2013 |
| 63 | *QYr.cbl-5D.1* | chr5D | 116776490 | 214072420 | Resistance | Yr_W056;MLM |  | NA | - |  |
| 64 | *QYr.cbl-5D.2* | chr5D | 126592669 | 142738088 | Susceptible | Yr_W034;FarmCPU |  | NA | - |  |
| 65 | *QYr.cbl-5D.3* | chr5D | 210441398 | 230295467 | Resistance | Yr_W031;MLM |  | NA | - |  |
| 66 | *QYr.cbl-5D.4* | chr5D | 464928096 | 472046355 | Resistance | Yr_W031;MLM |  | NA | - |  |
| 67 | *QYr.cbl-6A.1* | chr6A | 81804519 | 118784370 | Resistance | Yr_W031;MLM | - | *QYr.sicau-6A;*<br>*QYr.cim-6AL* | QRC-6A-II | Ma et al. 2019;<br>Pretorius et al. 2017 |
| 68 | *QYr.cbl-6A.2* | chr6A | 300689728 | 320565123 | Resistance | Yr_W031;MLM | - | NA | - |  |
| 69 | *QYr.cbl-6A.3* | chr6A | 547905103 | 571956512 | Resistance | Yr_W057;MLM,<br>Yr_W057;FarmCPU | - | *QYren.orz-6AL* | - | Vazquez et al. 2015 |
| 70 | QYr.cbl-6B.1 | chr6B | 465709752 | 482344304 | Resistance | Yr_W056;MLM | - | NA | - |  |
| 71 | QYr.cbl-6B.2 | chr6B | 555612474 | 576120483 | Resistance | Yr_W034;MLM,<br>Yr_W031;MLM | - | qNV.Yr-6B.4 | - | Jambuthenne et al. 2022 |
| 72 | QYr.cbl-6B.3 | chr6B | 684500621 | 690657692 | Resistance | Yr_W034;FarmCPU | - | NA | - |  |
| 73 | QYr.cbl-6B.4 | chr6B | 704466621 | 712363275 | Resistance | Yr_W057;MLM,<br>Yr_W031;MLM | - | qNV.Yr-6B.5 | - | Jambuthenne et al. 2022 |
| 74 | QYr.cbl-6D.1 | chr6D | 17237113 | 21496608 | Resistance | Yr_W056;FarmCPU | - | NA | - |  |
| 75 | QYr.cbl-6D.2 | chr6D | 439120610 | 441832675 | Resistance | Yr_W031;FarmCPU | - | NA | - |  |
| 76 | QYr.cbl-7A.1 | chr7A | 46731623 | 62964704 | Resistance | Yr_W034;MLM,<br>Yr_W034;FarmCPU | - | NA | - |  |
| 77 | QYr.cbl-7A.2 | chr7A | 122329203 | 154018812 | Resistance | Yr_W031;MLM,<br>Yr_W034;MLM | - | QYr.nwafu-7AS | - | Wu et al. 2020 |
| 78 | QYr.cbl-7A.3 | chr7A | 175432901 | 195276452 | Resistance | Yr_W034;MLM | - | NA | - |  |
| 79 | QYr.cbl-7A.4 | chr7A | 655547760 | 669698588 | Resistance | Yr_W031;FarmCPU | - | QYr-7AL.1 | - | Zhang et al. 2021 |
| 80 | QYr.cbl-7B.1 | chr7B | 601476179 | 616767883 | Resistance | Yr_W034;MLM | - | qNV.Yr-7B.3 | - | Jambuthenne et al. 2022 |
| 81 | QYr.cbl-7B.2 | chr7B | 632516145 | 632682787 | Resistance | Yr_W057;FarmCPU | - | NA | - |  |
| 82 | QYr.cbl-7D.1 | chr7D | 12875233 | 17840109 | Resistance | Yr_W031;MLM | - | NA | - |  |
| 83 | QYr.cbl-7D.2 | chr7D | 43114349 | 58790214 | Resistance | Yr_W031;MLM | Lr34/Yr18/Sr57 | Lr34/Yr18/Sr57;<br>QYr.YBZR-7DS;<br>QYr.hebau-7DS;<br>QYr.caas-7DS;<br>QYr.spa-7D;<br>QYr.AYH-7DS;<br>QYr.sgi-7D;<br>QYr.GX-7DS | QRC-7D-I | Krattinger et al. 2009;<br>Deng et al. 2022;<br>Yan et al. 2021;<br>Lu et al. 2009;<br>Bokore et al. 2017;<br>Long et al. 2021;<br>Ramburan et al. 2004; Wang et al. 2021 |
| 84 | QYr.cbl-7D.3 | chr7D | 156968435 | 159575148 | Resistance | Yr_W031;FarmCPU | - | NA | - |  |
| 85 | QYr.cbl-7D.4 | chr7D | 343246840 | 363165461 | Resistance | Yr_W031;MLM | - | NA | - |  |
| 86 | QYr.cbl-7D.5 | chr7D | 540945484 | 560901121 | Resistance | Yr_W034;MLM,<br>Yr_W056;MLM | - | NA | - |  |
| 87 | *QYr.cbl-7D.6* | chr7D | 614170347 | 634113034 | Resistance | Yr_W031;MLM,<br>Yr_W001;MLM | - | *QYrmx.nwafu-7DS*;<br>*QYr.nwafu-7DL* | - | Yuan et al. 2018;<br>Wu et al. 2020 |

Of the 87 loci detected, 81 were associated with reduced IT (resistance effect), while six were associated with increased IT (susceptibility effect) (**Table 2**). The susceptibility-associated loci were *QYr.cbl-1B.2* (chr1B: 39.7–54.1 Mb), *QYr.cbl-2D.4* (chr2D: 268.7–288.6 Mb), *QYr.cbl-4B.5* (chr4B: 113.3–133.2 Mb), *QYr.cbl-5A.6* (chr5A: 669.9–689.1 Mb), *QYr.cbl-5B.5* (chr5B: 510.5–523.2 Mb), and *QYr.cbl-5D.2* (chr5D: 126.6–142.7 Mb). All remaining loci contributed alleles associated with decreased disease severity.

QTL were unevenly distribution across the genome (**Table 2**). Chromosome 4B harbored the largest number of loci (10), followed by chromosome 5B (9) and chromosome 1B (8). Chromosomes 2D and 7D each contained multiple loci (7 and 6, respectively), while chromosomes 2A, 3A, 3B, 5A, 5D, 6A, 6B, 6D, 7A, and 7B also contributed distinct QTL.

A subset of loci was supported by more than one isolate, indicating genomic regions associated with resistance across different *Pst* isolates. In total, 23 QTL were detected for two or more isolates (**Table 2**). Among these, *QYr.cbl-3B.3* (chr3B: 619.3–676.4 Mb) showed the broad-spectrum resistance and was detected for W034, W031, W001, W057, and T022. *QYr.cbl-4B.8* (chr4B: 228.3–298.5 Mb) conferred resistance to W057, W056, W031, and T022. *QYr.cbl-5B.3* (chr5B: 92.4–296.9 Mb) was associated with W056, W031, and W034. Similarly, *QYr.cbl-2D.5* (chr2D: 476.0–507.6 Mb) was detected for W057, W031, and W056, and *QYr.cbl-1B.6* (chr1B: 510.2–563.5 Mb) was associated with W056, W031, and W001. Across isolates, W031 and W056 contributed to the greatest number of shared loci, whereas W001 and T022 were involved in fewer multi-isolate regions. These shared loci indicate partial overlap in the genetic basis of resistance among isolates, while numerous isolate-specific QTL were also identified.

Several loci were detected under both MLM and FarmCPU models, providing additional statistical support for these associations (**Table 2**). Fourteen QTL showed multi-model support, either within the same isolate or across isolates. Examples include *QYr.cbl-1B.1* (W056 detected under both MLM and FarmCPU), *QYr.cbl-1B.6* (W056 under MLM and FarmCPU and W001 under FarmCPU), *QYr.cbl-2A.1* (W057 under both models), *QYr.cbl-2D.5* (W031 under MLM and FarmCPU), *QYr.cbl-3B.3* (W031 and W001 detected under both models), *QYr.cbl-4A.2* (W056 and W001 under both models), *QYr.cbl-4B.8* (W057 and W056 under both models), *QYr.cbl-5A.1* (W056 and W034 under different models), *QYr.cbl-5B.3* (W031 under both MLM and FarmCPU), *QYr.cbl-6A.3* (W057 under MLM and FarmCPU), and *QYr.cbl-7A.1* (W034 under both models). The detection of these loci across statistical models or *Pst* isolates increases confidence in their robustness.

Additionally, putative candidate genes located within each of the 87 LD-defined QTL intervals were catalogued to assess gene density and refine priorities for downstream analyses (**Table S5**). The number of annotated genes per QTL varied substantially, reflecting differences in interval size and local genomic structure. Gene content ranged from as few as four genes in *QYr.cbl-7B.2* and seven genes in *QYr.cbl-7D.3* to as many as 817 genes in *QYr.cbl-5B*.*3*, which represented the most gene-dense interval identified in this study. Other large intervals included *QYr.cbl-4A.2* (539 genes), *QYr.cbl-2D.1* (489 genes), *QYr.cbl-1B.6* (433 genes), and *QYr.cbl-3B.3* (424 genes). For the majority of loci, gene counts fell within an intermediate range, typically between ∼50 and 300 genes, indicating moderate interval sizes suitable for further refinement (**Table S5**). These gene inventories provide a foundation for subsequent candidate gene prioritization based on functional annotation, expression profiling, and haplotype analysis.

### Co-localization with known Yr genes, Yr QTL and QTL-rich clusters (QRCs)

Comparative analysis of the 87 LD-defined QTL intervals with the physical positions of designated or cloned *Yr* genes identified co-localization for ten loci distributed across five chromosomes (**Table 2**). On chromosome 1B, *QYr.cbl-1B.1* overlapped with *Yr84* (Klymiuk et al. 2022), while *QYr.cbl-1B.2* and *QYr.cbl-1B.3* mapped within the reported interval of *Yr85* (Feng et al. 2023). On chromosome 2D, *QYr.cbl-2D.1* co-localized with *Yrq1* in the distal region of 2DS (Cao et al. 2012). On chromosome 3D, *QYr.cbl-3D.2* overlapped with *Yr71* (Bariana et al. 2016). Chromosome 4A contained *QYr.cbl-4A.2* within the interval reported for *Yr60* (Herrera-Foessel et al. 2015). A major region on chromosome 4B, *QYr.cbl-4B.8*, co-localized with *Yr62* and *Yr50* (Liu et al. 2013; Lu et al. 2014), while *QYr.cbl-4B.10* corresponded to the reported position of *Yr68* (Wang et al. 2021). On chromosome 5A, *QYr.cbl-5A.5* aligned with *Yr34* (Bariana et al. 2006). Finally, *QYr.cbl-7D.2* overlapped with the *Yr18* on chromosome 7DS (Krattinger et al. 2009).

In addition to cloned genes, 34 of the 87 QTLs co-localized with at least one previously reported stripe rust QTL (**Table 2**). On chromosome 1B, three loci showed overlap with previously reported QTL: *QYr.cbl-1B.1* co-localized with *QYr.cim-1BS*, *QYr.hbaas-1BS*, and *QYr.nwafu-1BS.1* (Jia et al. 2020; Wu et al. 2020; Zhang et al. 2022); *QYr.cbl-1B.6* corresponded to *QYrpd.swust-1BL* (Zhou et al. 2022); and *QYr.cbl-1B.7* overlapped *QYr.nmbu.1B* (Lin et al. 2023). On chromosome 2A, *QYr.cbl-2A.5* and *QYr.cbl-2A.6* overlapped *QYr.spa-2A* loci and *QYr.lrdc-2A.2* (Bokore et al. 2017; Farzand et al. 2022), while on chromosome 2D, *QYr.cbl-2D.1* and *QYr.cbl-2D.3* corresponded to *QYr.nwafu-2DS* (Wu et al. 2020) and *QYr.cbl-2D.5* overlapped *QYr.ufs-2D* (Agenbag et al. 2012). On chromosome 4B, *QYr.cbl-4B.4* overlapped *QYrva.vt-4BL* (Christopher et al. 2013) and *QYr.cbl-4B*.*7* corresponded to *QYrhm.nwafu-4B/6A* (Yuan et al. 2018), while *QYr.cbl-4D.2* aligned with *QYr.hbaas-4DL* (Jia et al. 2020). The largest number of overlaps occurred on chromosome 5B, where *QYr.cbl-5B.1*, *QYr.cbl-5B.3*, *QYr.cbl-5B.6*, *QYr.cbl-5B.8*, and *QYr.cbl-5B.9* each co-localized with previously reported 5B QTL (**Table 2**). Additionally, three overlapping QTLs were identified on 5A, two overlaps for each on 3A, 6A, 6B, 7A, 7D and one overlap on 3D, 7B chromosome.

Fifteen of the 87 QTL mapped within previously defined QRC (**Table 2**) (Tong et al. 2024). On chromosome 1B, *QYr.cbl-1B.1* fell within *QRC-1B-I* and *QYr.cbl-1B.6* within *QRC-1B-IV.* On chromosome 2A, *QYr.cbl-2A.5* mapped within *QRC-2A-V*, and on chromosome 2D, *QYr.cbl-2D.1* corresponded to *QRC-2D-I*. On chromosome 3A, *QYr.cbl-3A.1* fell within *QRC-3A-II*. On chromosome 4B, *QYr.cbl-4B.8* was located within *QRC-4B-I*. On chromosome 4D, *QYr.cbl-4D.2* corresponded to *QRC-4D-II*. On chromosome 5A, both *QYr.cbl-5A.5* and *QYr.cbl-5A.6* mapped within *QRC-5A-III*. On chromosome 5B, *QYr.cbl-5B.1* fell within *QRC-5B-I*, *QYr.cbl-5B.8* within *QRC-5B-V*, and *QYr.cbl-5B.9* within *QRC-5B-VII*. On chromosome 6A, *QYr.cbl-6A.1* mapped within *QRC-6A-II*. Finally, on chromosome 7D, *QYr.cbl-7D.2* corresponded to *QRC-7D-I*. The localization of these 15 QTL within established resistance-dense regions indicates partial overlap with previously recognized genomic hotspots.

### Novel resistance loci

A substantial proportion (n=46) of the mapped loci identified in this study did not overlap with previously reported *Yr* genes or QTL and therefore represent potentially novel regions associated with stripe rust resistance (**Table 2**). While many of these loci were detected for individual isolates, a subset (n=8) showed stronger resistance across isolates or models. For example, *QYr.cbl-3B.3* was detected across five isolates (W034, W031, W001, W057, and T022) and by both MLM and FarmCPU models, representing the most consistently supported novel region in the study. Similarly, *QYr.cbl-4B.2* was detected across multiple isolates (W031, W056, and W057) and across both models, while *QYr.cbl-2A.1* was detected by two isolates and both MLM and FarmCPU. In addition, *QYr.cbl-7A.1* was detected for isolate W034 under both MLM and FarmCPU models, providing independent model support for this locus despite its isolate-specific detection. Additional loci such as *QYr.cbl-1D.1*, *QYr.cbl-2A.2*, and *QYr.cbl-7D.5* were detected across multiple isolates. Collectively, these findings highlight the extensive unexplored resistance diversity present within the Watkins panel and identify several genomic regions that warrant further investigation through fine mapping and functional validation.

### Haplotype analysis

A subset of 29 QTL identified by multiple stripe rust isolates, detected by both GWAS models, and/or co-localizing with previously designated *Yr* genes were considered high-confidence loci and selected for haplotype analysis (**Table S6**). Significant multi-SNP haplotypes were identified for all loci except *QYr.cbl-5B.2* and *QYr.cbl-7D.6* (**Fig. 5; Figs. S6–S9**). The size of haplotype blocks varied substantially among QTL, ranging from two SNPs in *QYr.cbl-7D.2*, three SNPs in *QYr.cbl-4B.7*, *QYr.cbl-4B.9*, *QYr.cbl-5A.1*, and *QYr.cbl-5B.3* to 47 SNPs in *QYr.cbl-4B.2*, followed by 25 SNPs in *QYr.cbl-6A.3* and 23 SNPs in *QYr.cbl-7A.1* (**Table S8-S36**). Across 29 loci, the average haplotype block contained approximately 10 SNPs per QTL. The number and frequency of haplotypes also varied considerably among loci and were not necessarily correlated with the number of SNPs within the haplotype block. For example, although *QYr.cbl-6A.3* contained a haplotype block of 25 SNPs (**Table S29**), these markers resolved into only two haplotypes, one associated with susceptibility and the other with resistance. A similar pattern was observed for *QYr.cbl-4A.2* (**Table S18**), where 15 SNPs formed three haplotypes, and for *QYr.cbl-4B.2* (**Table S19**), where 47 SNPs also resolved into only three haplotypes. In contrast, a greater diversity of haplotypes was detected for some loci, with the highest number observed for *QYr.cbl-1D.1* (n = 13) (**Table S11**), followed by *QYr.cbl-4B.10* with 12 haplotypes (**Table S24**). Notably, even at loci with multiple haplotypes, only one and occasionally two haplotypes were consistently associated with significantly reduced stripe rust infection scores. Collectively, these results refine the GWAS signals into discrete haplotype blocks and identify favorable allelic combinations that can serve as diagnostic markers for marker-assisted selection and pyramiding of stripe rust resistance in wheat breeding programs.

**Figure 5.**
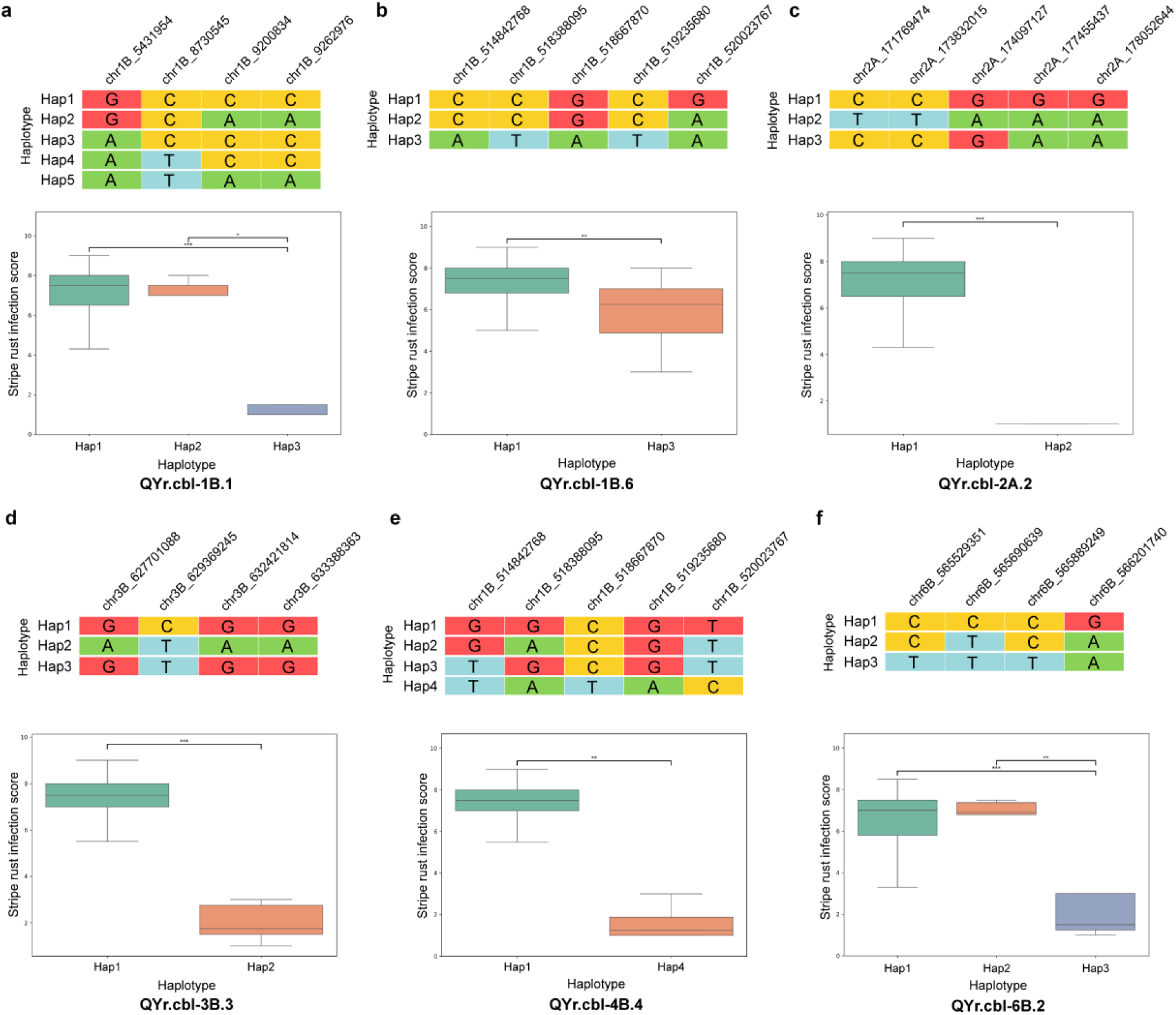
Haplotype effects at selected stripe rust resistance QTL. Haplotype composition (top) and phenotypic effects (bottom) are shown for (a) *QYr.cbl-1B.1*, (b) *QYr.cbl-1B.6*, (c) *QYr.cbl-2A.2*, (d) *QYr.cbl-3B.3*, (e) *QYr.cbl-4B.4*, and (f) *QYr.cbl-6B.2*. Colored cells indicate SNP alleles (A, T, G, C) within each LD-defined QTL interval. Boxplots show stripe rust infection scores (0–9) for accessions carrying each haplotype. Asterisks indicate significant differences between haplotypes based on pairwise Mann–Whitney U tests (*P* < 0.05, *P* < 0.01, *P* < 0.001).

### Watkin accessions carrying multiple resistance loci

A subset of Watkins landraces carried favorable alleles at multiple high-confidence QTL (**Table S7**). WATDE0042, WATDE0076, and WATDE0776 each carried favorable alleles at 17 QTL, including loci aligned with known resistance genes *Yr18*, *Yr68, Yr71,* and *Yr85*, together with several novel loci (e.g., *QYr.cbl-1D.1, QYr.cbl-2A.1, QYr.cbl-3B.3*, and *QYr.cbl-7A.1*). Similarly, WATDE0694, WATDE0581, and WATDE0895 carried favorable alleles at 12–14 loci, combining multiple known resistance regions including *Yr84, Yrq1, Yr60/Yr51*, and *Yr62/Yr50* with additional QTL detected in the present study. These accessions therefore represent particularly valuable genetic resources for resistance gene discovery and for breeding strategies aimed at pyramiding multiple resistance loci.

## Discussion

Landrace collections such as Watkins, especially when paired with full resequencing data, offer an opportunity to reintroduce lost variation and diversify resistance sources before further breakdown events occur (Wingen et al. 2017; Cheng et al. 2024). Watkins landraces are a proven resource of stripe rust resistance. Over the past decade, several Yr genes have been identified from individual Watkins accessions, including *Yr47, Yr51, Yr57, Yr60, Yr63, Yr72, Yr80, Yr81,* and *Yr82* (Bansal et al. 2011; Bariana et al. 2014; Chhetri et al. 2023; Gessese et al. 2019; Mackenzie et al. 2023; Nsabiyera et al. 2018; Pakeerathan et al. 2019; Randhawa et al. 2014; Randhawa et al. 2015). In each case, a single resistant accession was crossed to a susceptible parent, and the locus was mapped in a biparental population. While powerful, this approach sampled only one resistance source at a time. Consequently, the broader allelic diversity and multi-locus architecture present across the Watkins panel remained largely unexplored.

A few studies examined larger portions of the collection using association scans or genomic prediction (Bansal et al. 2013; Daetwyler et al. 2014; Pasam et al. 2017). However, these efforts relied on lower-density marker systems and lacked the genomic resolution needed to fully capture LD structure, rare alleles, and complex haplotypes. The whole-genome resequencing dataset generated by Cheng et al. (2024) represents a major advance, providing the marker density required to dissect resistance at high resolution. To our knowledge, this is the first study, beyond the original sequencing report, that integrates the Watkins high-density resequencing dataset with phenotyping against six diverse *Pst* isolates, offering a comprehensive view of stripe rust resistance within the panel.

Ten loci detected in this study co-localized with physically mapped or cloned *Yr* genes. The recovery of these regions supports both the GWAS framework and the biological relevance of these loci. Importantly, *Yr60* was originally characterized from a Watkins accession (Bariana et al. 2014), and *Lr34/Yr18/Sr57* is known to occur within Watkins diversity (Wamalwa et al. 2019). Their detection here confirms that favorable haplotypes at these regions remain present and detectable at the population scale. However, co-localization does not prove gene identity. Because these signals are defined by LD-based physical overlap, some may represent tightly linked but distinct loci rather than the canonical designated or cloned *Yr* genes. Haplotype comparison, diagnostic marker validation, fine mapping, and functional assays will be necessary to determine whether these favorable alleles represent identical genes, alternative alleles, or novel functional variants.

From a modeling perspective, the contrast between GLM, MLM, and FarmCPU highlights the importance of appropriate statistical control. GLM showed clear inflation, confirming that simple models are inadequate in structured landrace panels. MLM and FarmCPU both controlled-inflation effectively but differ in sensitivity. We retained loci detected by either MLM or FarmCPU because moderate-effect loci may be captured by one framework but not the other. Restricting interpretation only to loci shared across models, risks discarding biologically meaningful signals (Pasam et al. 2017). At the same time, loci supported by both models were considered especially robust.

A key practical outcome of this work is the identification of Watkins accessions carrying favorable alleles at multiple *Yr*-aligned loci. Such multi-locus backgrounds represent strong candidates for pyramiding and pre-breeding. However, marker validation and haplotype comparison with known donor lines should precede deployment. Phenotyping was conducted under controlled conditions at the seedling stage; therefore, we have only mapped all-stage resistance (ASR) loci and field phenotyping could help identify adult-plant resistance (APR) loci. However, field phenotyping of Watkins accessions is quite challenging due to different growth habits and phenology of accessions. In our study, only 297 of the 827 Watkins accessions were evaluated, indicating that a substantial portion of the genetic diversity within the collection remains unexplored. Moreover, certain *Yr* genes previously reported in Watkins landraces were not detected in the GWAS. For example, *Yr47*, reported in accession WATDE0042 (Bansal et al. 2011), was not identified by either the MLM or FarmCPU models, highlighting the inherent limitations of association mapping and the possibility that some resistance loci may remain undetected due to allele frequency or population structure.

In summary, the Watkins collection harbors extensive and structured variation for stripe rust resistance, spanning both known resistance regions and potentially novel loci. By leveraging high-density resequencing and multi-isolate phenotyping, this study provides a broader view of resistance architecture. These findings lay the groundwork for systematic validation and strategic deployment of landrace-derived alleles to strengthen the genetic resilience of modern wheat.

## Acknowledgements

The authors acknowledge the support and help received from members of the Cereal Breeding Lab (CBL).

## Funding

The funding for this research was provided by Saskatchewan Ministry of Agriculture (Project ID: ADF20200205), Saskatchewan Wheat Development Commission (Project ID: 173-201127), Alberta Grains (Project ID: 2021AWC100B), and Manitoba Crop Alliance (Project ID: MCA2184).

## Author contribution statement

GSB conceived the idea and supervised the project. GSB performed the stripe rust phenotyping. JS and MJAA performed bioinformatic and other data analyses. JS and GSB wrote the manuscript. JS, MJAA, and NK prepared figures and tables. SH and RSK made significant intellectual contributions. All authors reviewed and approved the final manuscript.

## Supplementary Figures and Table

**Figure S1.**
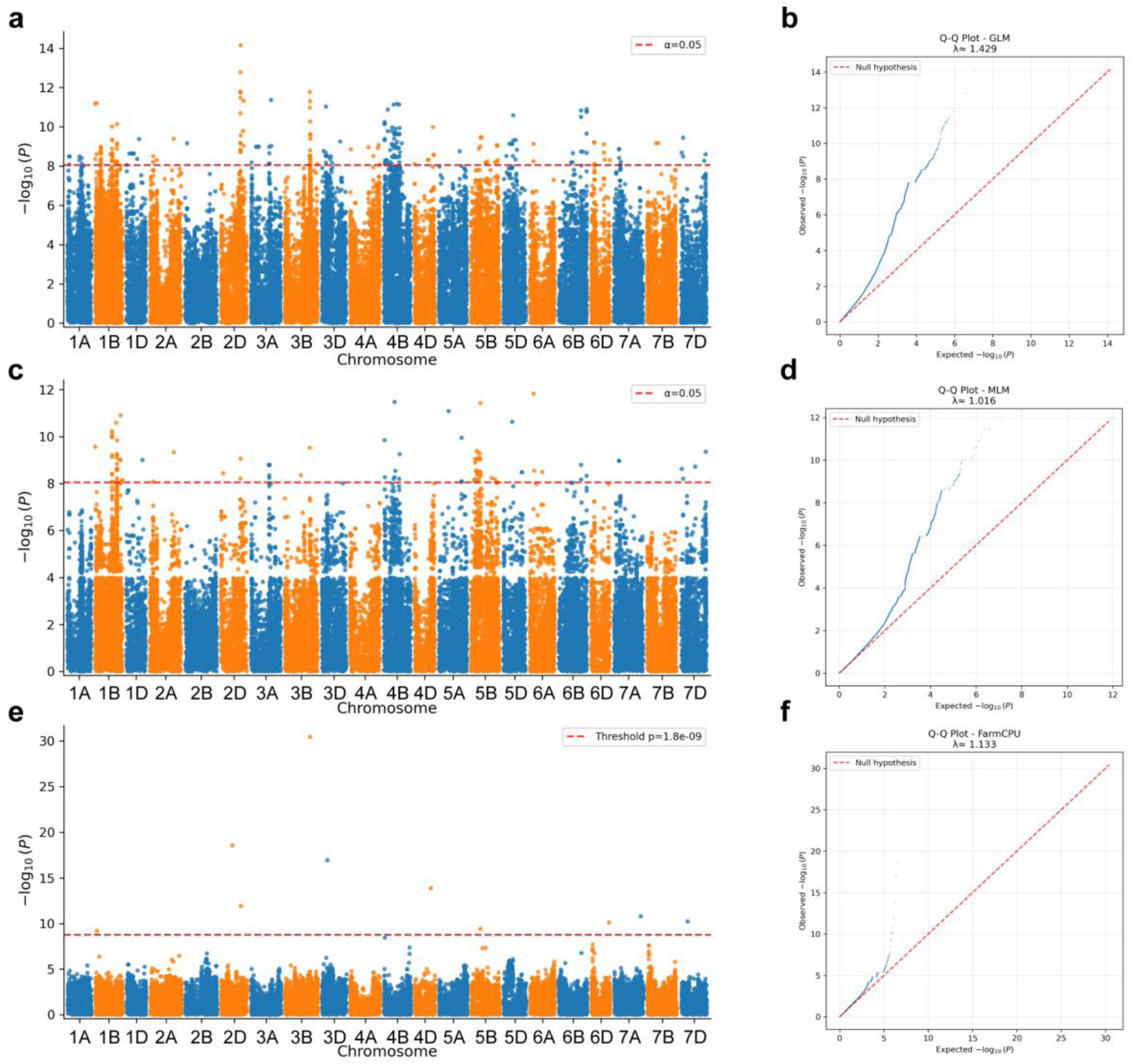
Genome-wide association analysis for stripe rust isolate W031 using GLM, MLM, and FarmCPU models. (a) Manhattan plot from the GLM analysis showing genome-wide marker–trait associations across the 21 wheat chromosomes. The red dashed line indicates the significance threshold (α = 0.05). (b) Q–Q plot for the GLM model illustrating the deviation of observed from expected –log₁₀(P) values, with an inflation factor (λ) of 1.429. (c) Manhattan plot from the MLM analysis incorporating population structure and kinship. The red dashed line denotes the significance threshold (α = 0.05). (d) Q–Q plot for the MLM model (λ = 1.016), indicating improved control of false positives relative to GLM. (e) Manhattan plot from the FarmCPU analysis, with the genome-wide significance threshold (P = 1.8 × 10⁻⁹) shown by the red dashed line. (f) Q–Q plot for the FarmCPU model (λ = 1.133), demonstrating effective control of population structure and cryptic relatedness while retaining significant associations.

**Figure S2.**
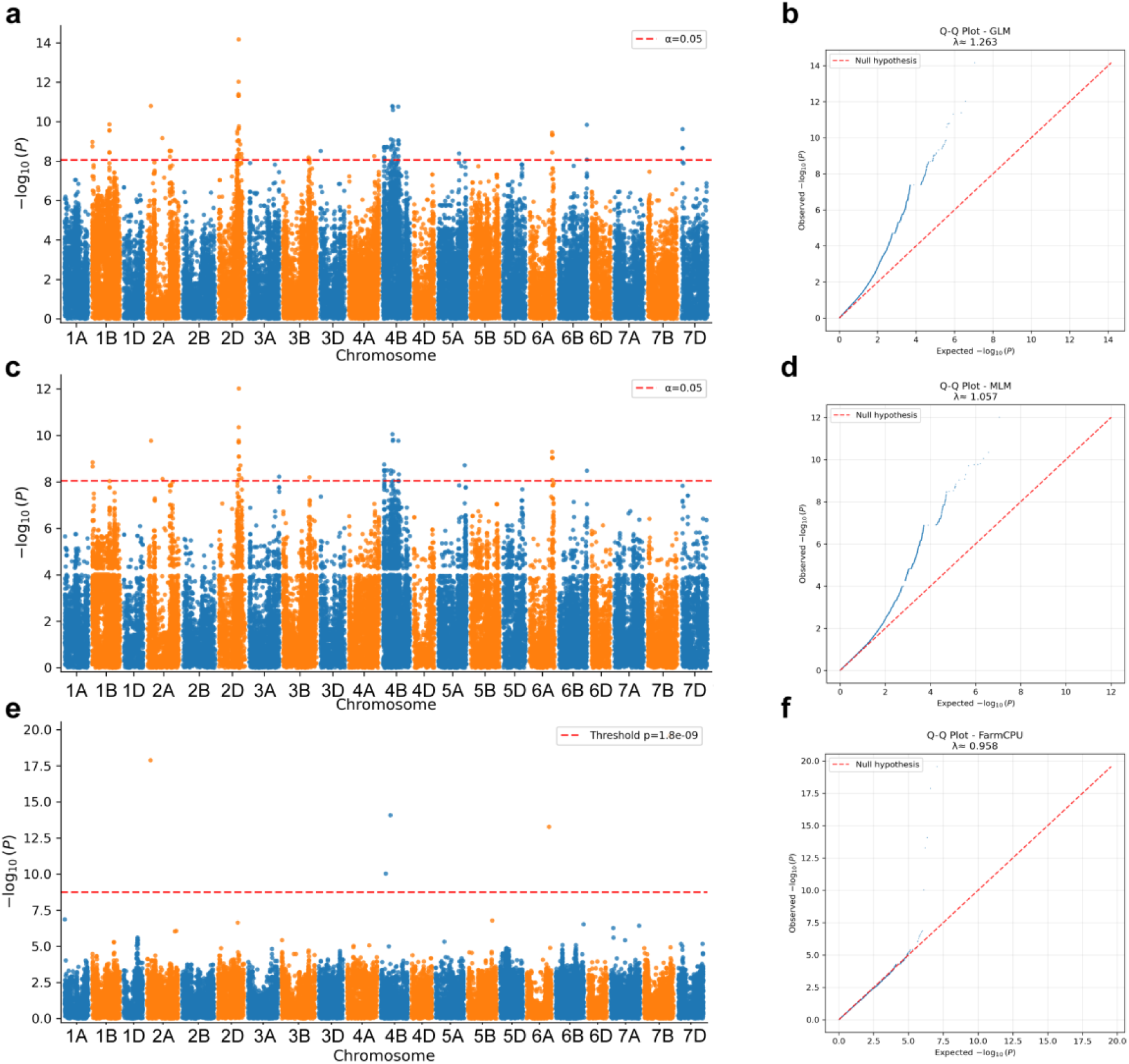
Genome-wide association analysis for stripe rust isolate W057 using GLM, MLM, and FarmCPU models. (a) Manhattan plot from the GLM analysis showing genome-wide marker–trait associations across the 21 wheat chromosomes. The red dashed line indicates the significance threshold (α = 0.05). (b) Q–Q plot for the GLM model illustrating the deviation of observed from expected –log₁₀(P) values, with an inflation factor (λ) of 1.263. (c) Manhattan plot from the MLM analysis incorporating population structure and kinship. The red dashed line denotes the significance threshold (α = 0.05). (d) Q–Q plot for the MLM model (λ = 1.057), indicating improved control of false positives relative to GLM. (e) Manhattan plot from the FarmCPU analysis, with the genome-wide significance threshold (P = 1.8 × 10⁻⁹) shown by the red dashed line. (f) Q–Q plot for the FarmCPU model (λ = 0.958), demonstrating effective control of population structure and cryptic relatedness while retaining significant associations.

**Figure S3.**
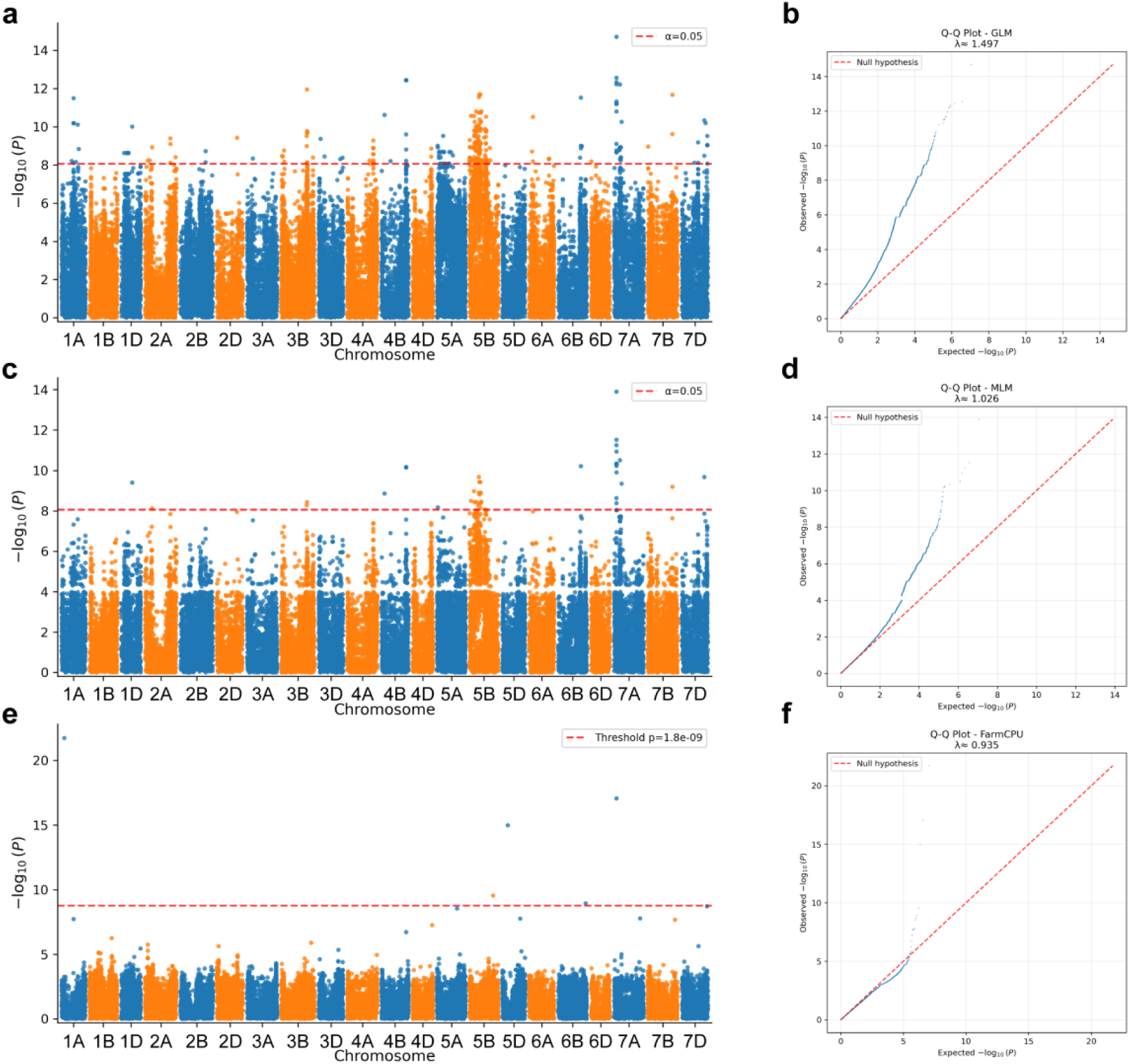
Genome-wide association analysis for stripe rust isolate W034 using GLM, MLM, and FarmCPU models. (a) Manhattan plot from the GLM analysis showing genome-wide marker–trait associations across the 21 wheat chromosomes. The red dashed line indicates the significance threshold (α = 0.05). (b) Q–Q plot for the GLM model illustrating the deviation of observed from expected –log₁₀(P) values, with an inflation factor (λ) of 1.497. (c) Manhattan plot from the MLM analysis incorporating population structure and kinship. The red dashed line denotes the significance threshold (α = 0.05). (d) Q–Q plot for the MLM model (λ = 1.026), indicating improved control of false positives relative to GLM. (e) Manhattan plot from the FarmCPU analysis, with the genome-wide significance threshold (P = 1.8 × 10⁻⁹) shown by the red dashed line. (f) Q–Q plot for the FarmCPU model (λ = 0.935), demonstrating effective control of population structure and cryptic relatedness while retaining significant associations.

**Figure S4.**
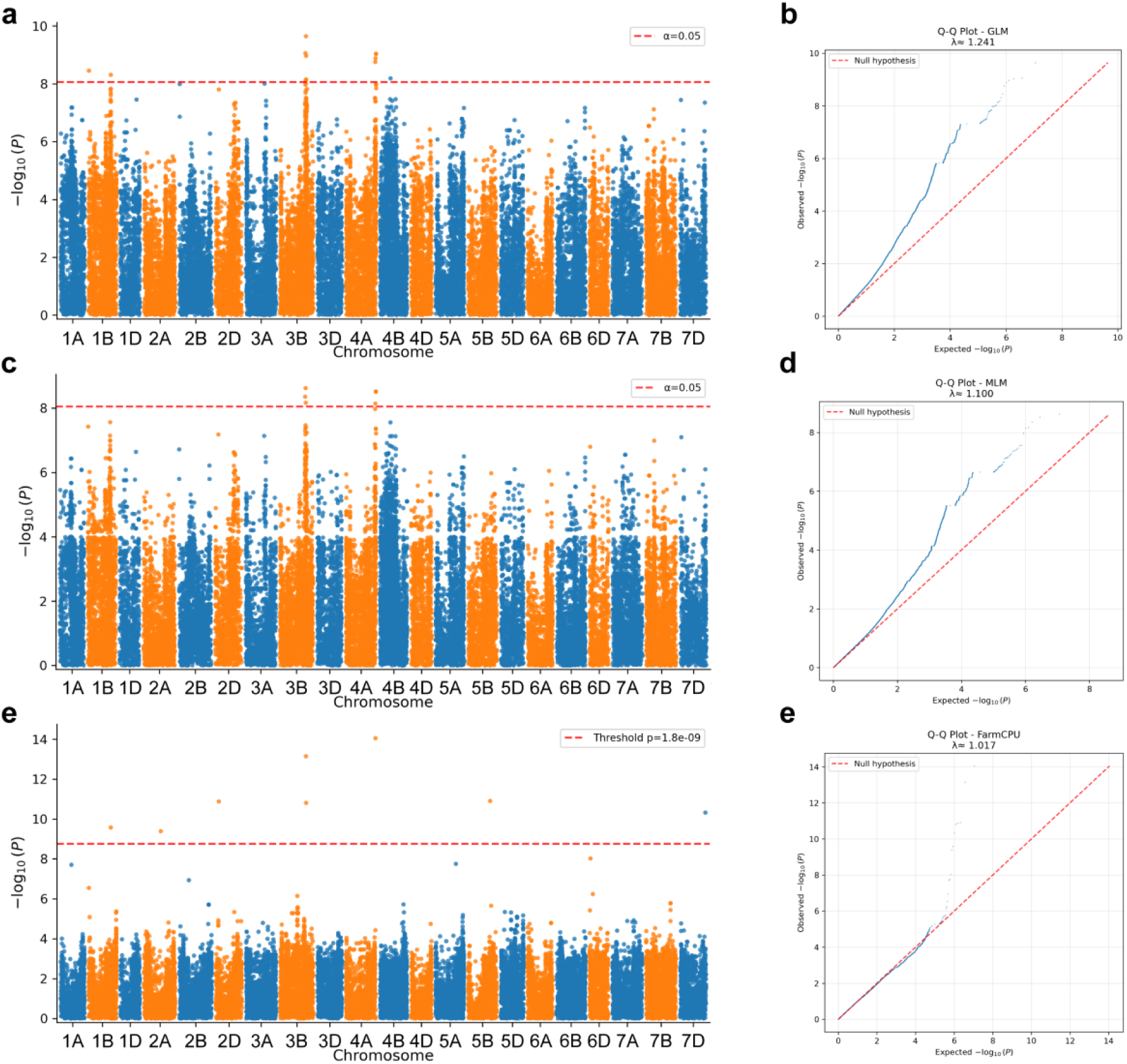
Genome-wide association analysis for stripe rust isolate W001 using GLM, MLM, and FarmCPU models. (a) Manhattan plot from the GLM analysis showing genome-wide marker–trait associations across the 21 wheat chromosomes. The red dashed line indicates the significance threshold (α = 0.05). (b) Q–Q plot for the GLM model illustrating the deviation of observed from expected –log₁₀(P) values, with an inflation factor (λ) of 1.241. (c) Manhattan plot from the MLM analysis incorporating population structure and kinship. The red dashed line denotes the significance threshold (α = 0.05). (d) Q–Q plot for the MLM model (λ = 1.100), indicating improved control of false positives relative to GLM. (e) Manhattan plot from the FarmCPU analysis, with the genome-wide significance threshold (P = 1.8 × 10⁻⁹) shown by the red dashed line. (f) Q–Q plot for the FarmCPU model (λ = 1.017), demonstrating effective control of population structure and cryptic relatedness while retaining significant associations.

**Figure S5.**
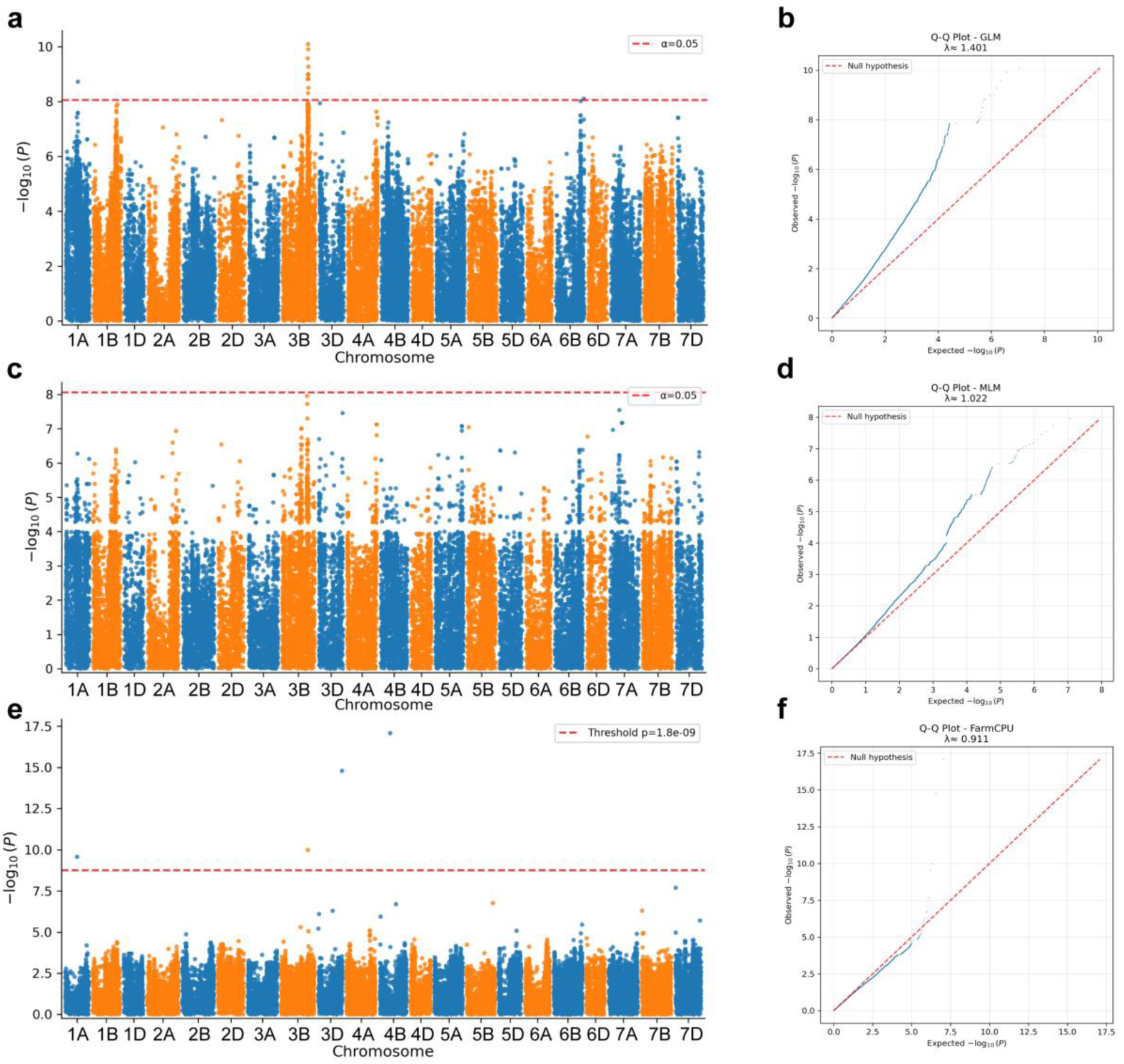
Genome-wide association analysis for stripe rust isolate T022 using GLM, MLM, and FarmCPU models. (a) Manhattan plot from the GLM analysis showing genome-wide marker–trait associations across the 21 wheat chromosomes. The red dashed line indicates the significance threshold (α = 1.401). (b) Q–Q plot for the GLM model illustrating the deviation of observed from expected –log₁₀(P) values, with an inflation factor (λ) of 1.401. (c) Manhattan plot from the MLM analysis incorporating population structure and kinship. The red dashed line denotes the significance threshold (α = 0.05). (d) Q–Q plot for the MLM model (λ = 1.022), indicating improved control of false positives relative to GLM. (e) Manhattan plot from the FarmCPU analysis, with the genome-wide significance threshold (P = 1.8 × 10⁻⁹) shown by the red dashed line. (f) Q–Q plot for the FarmCPU model (λ = 0.911), demonstrating effective control of population structure and cryptic relatedness while retaining significant associations.

**Figure S6.**
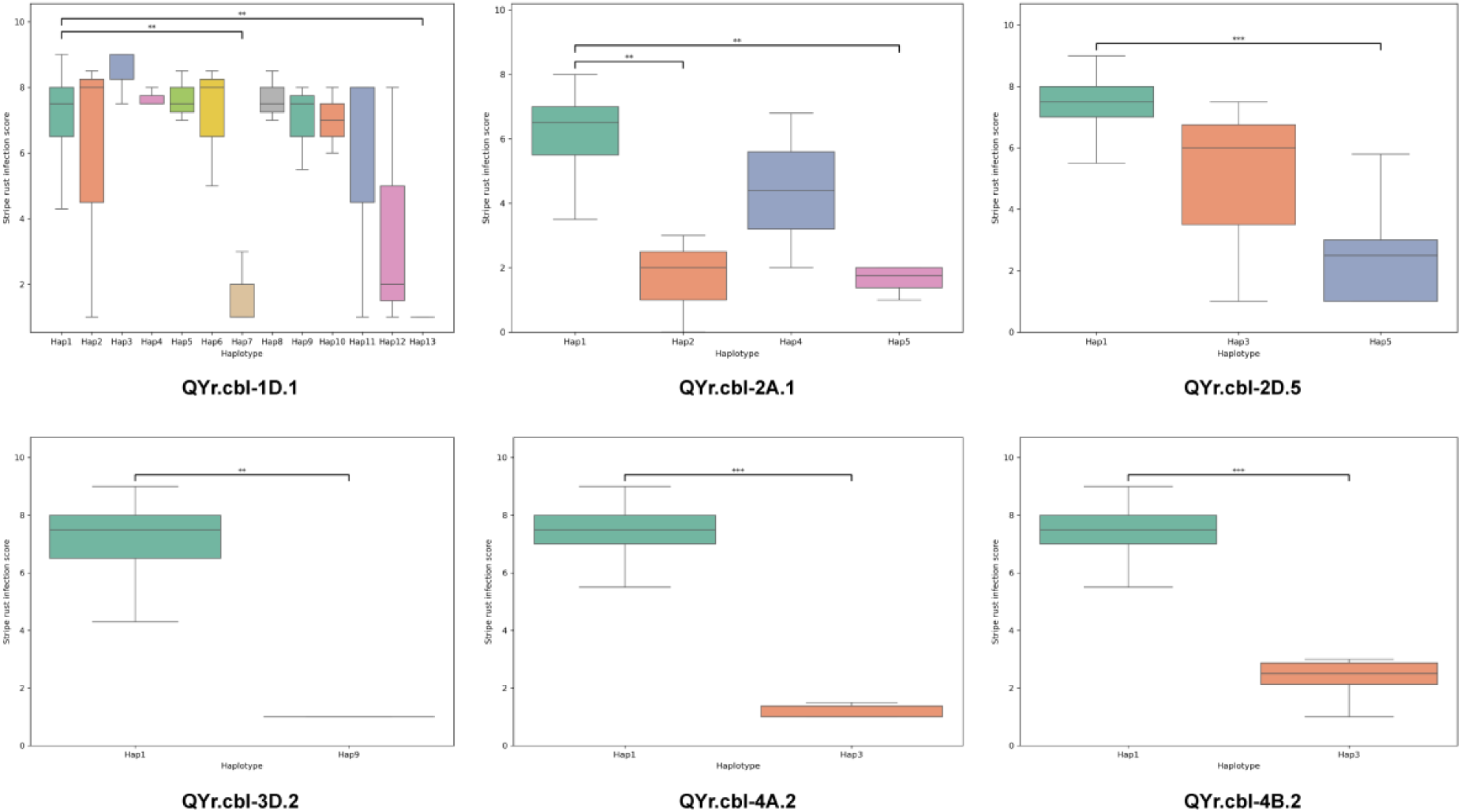
Haplotype effects of stripe rust resistance QTL identified in the Watkins panel. Boxplots show stripe rust infection scores (0–9) for accessions carrying different haplotypes at *QYr.cbl-1D.1*, *QYr.cbl-2A.1*, *QYr.cbl-2D.5*, *QYr.cbl-3D.2*, *QYr.cbl-4A.2*, and *QYr.cbl-4B.2*. Boxes represent the interquartile range with median values shown by horizontal lines. Asterisks indicate significant differences between haplotypes based on pairwise Mann–Whitney U tests (*P* < 0.01, *P* < 0.001). Resistant haplotypes are associated with lower infection scores.

**Figure S7.**
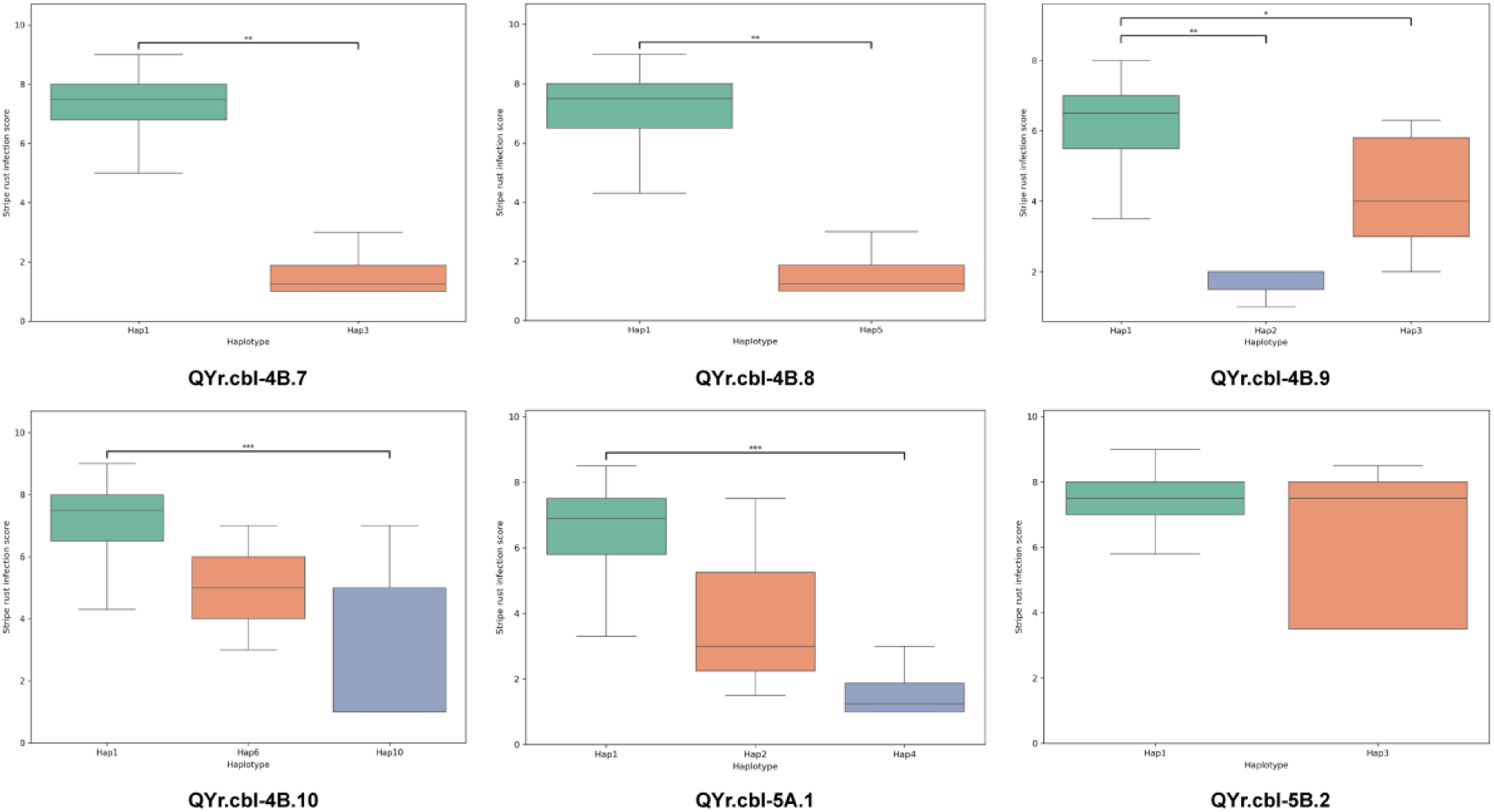
Haplotype effects of stripe rust resistance QTL identified in the Watkins panel. Boxplots show stripe rust infection scores (0–9) for accessions carrying different haplotypes at *QYr.cbl-4B.7, QYr.cbl-4B.8, QYr.cbl-4B.9, QYr.cbl-4B.10, QYr.cbl-5A.1,* and *QYr.cbl-5B.2.* Boxes represent the interquartile range with median values indicated by horizontal lines. Asterisks denote significant differences between haplotypes based on pairwise Mann–Whitney U tests (*P* < 0.05, *P* < 0.01, *P* < 0.001), with resistant haplotypes showing lower infection scores.

**Figure S8.**
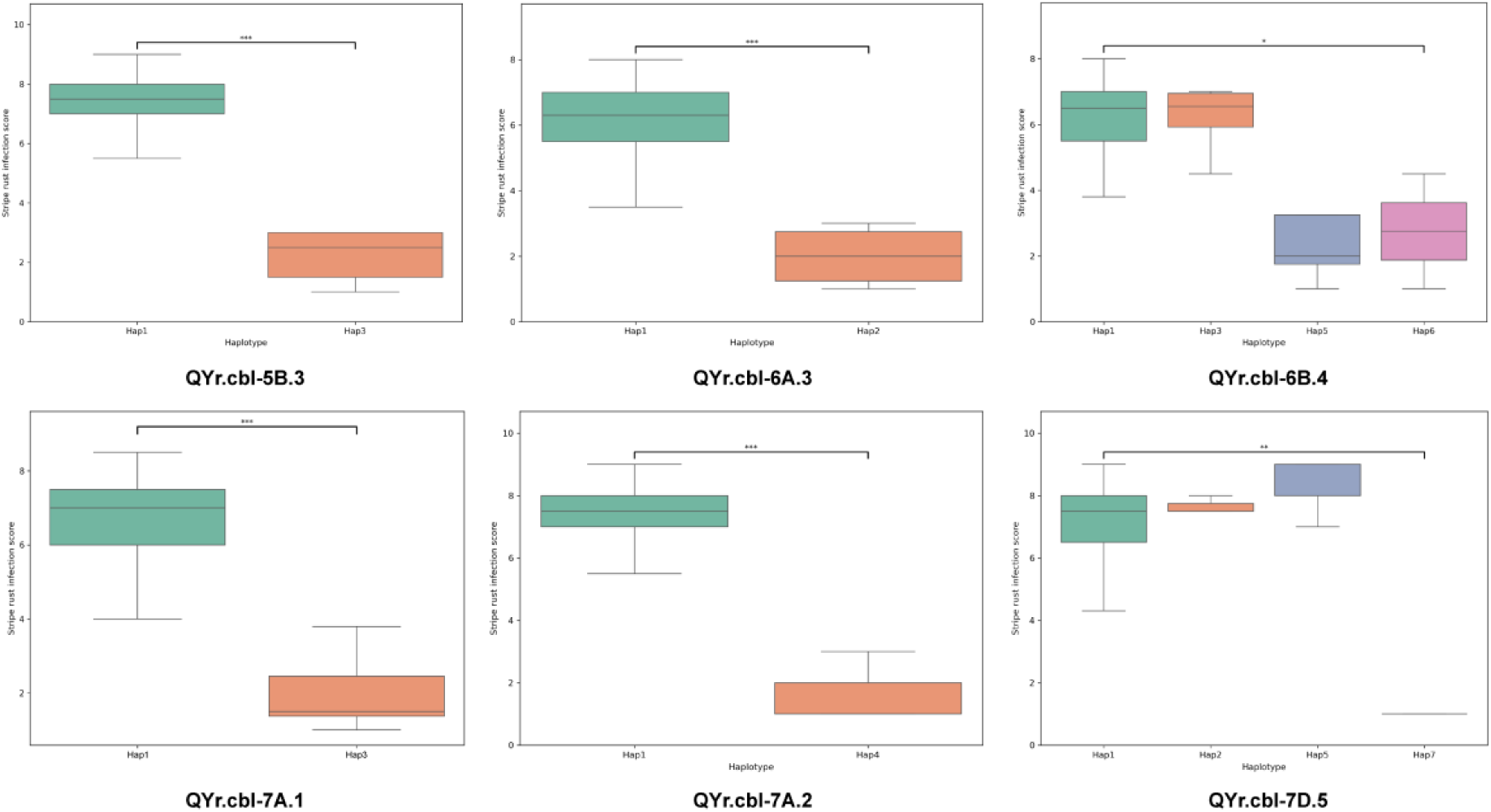
Haplotype effects of stripe rust resistance QTL identified in the Watkins panel. Boxplots show stripe rust infection scores (0–9) for accessions carrying different haplotypes at *QYr.cbl-5B.3, QYr.cbl-6A.3, QYr.cbl-6B.4, QYr.cbl-7A.1, QYr.cbl-7A.2,* and *QYr.cbl-7D.5.* Boxes represent the interquartile range with median values indicated by horizontal lines. Asterisks denote significant differences between haplotypes based on pairwise Mann–Whitney U tests (*P* < 0.05, *P* < 0.01, *P* < 0.001), with resistant haplotypes associated with lower infection scores.

**Figure S9.**
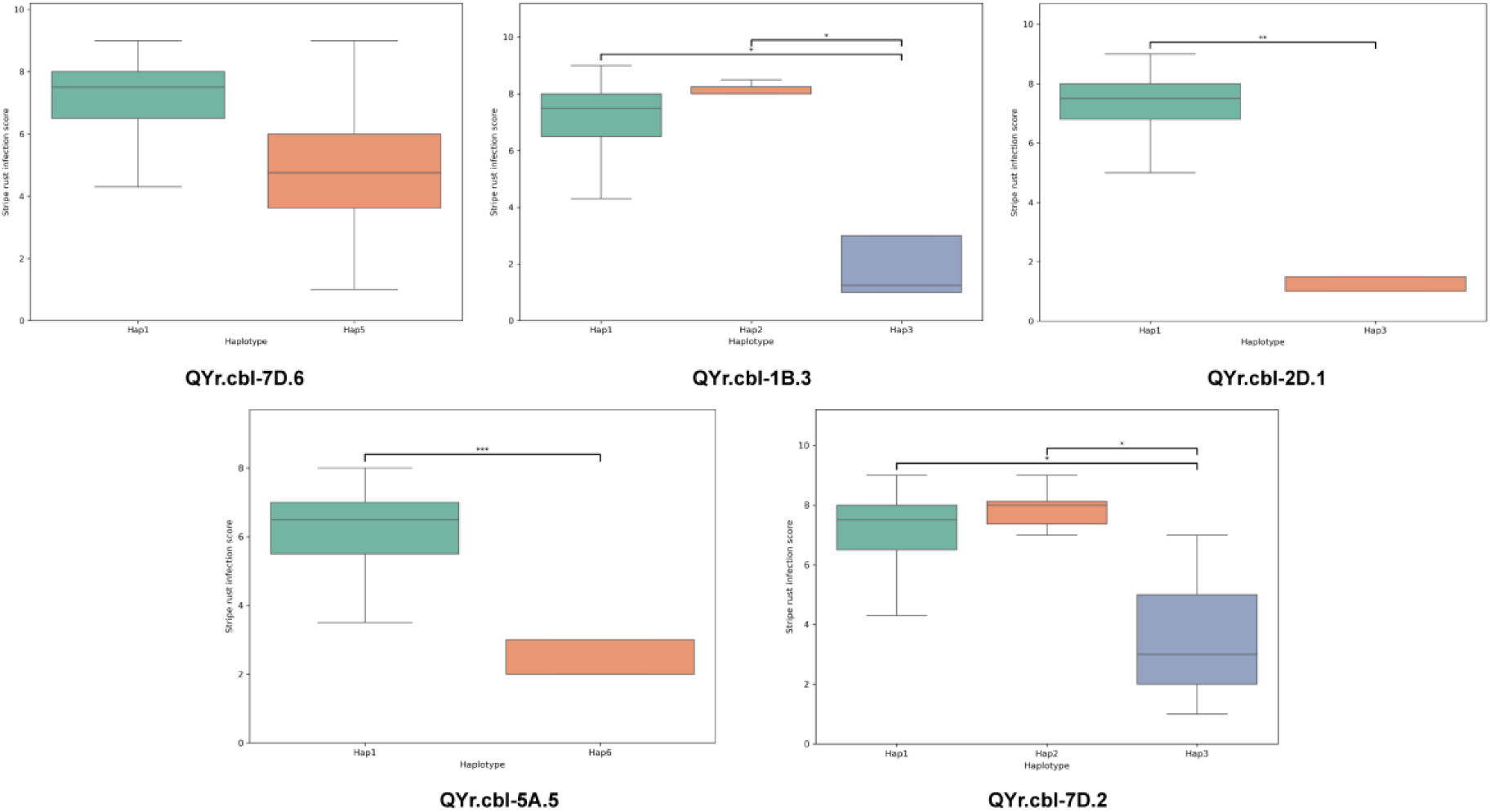
Haplotype effects of stripe rust resistance QTL identified in the Watkins panel. Boxplots show stripe rust infection scores (0–9) for accessions carrying different haplotypes at *QYr.cbl-7D.6, QYr.cbl-1B.3, QYr.cbl-2D.1, QYr.cbl-5A.5,* and *QYr.cbl-7D.2*. Boxes represent the interquartile range with median values indicated by horizontal lines. Asterisks denote significant differences between haplotypes based on pairwise Mann–Whitney U tests (*P* < 0.05, *P* < 0.01, *P* < 0.001), with resistant haplotypes associated with lower infection scores.

## Supplementary Tables

**Table S1.** Phenotypic responses of Watkins wheat landraces to six diverse stripe rust isolates.

**Table S2.** Genome-wide SNP marker counts across 21 wheat chromosomes at raw, quality-filtered, and LD-pruned stages.

**Table S3.** Genome-wide significant quantitative trait nucleotides (QTNs) identified across six stripe rust isolates using MLM and FarmCPU

**Table S4.** Detailed characteristics of representative stripe rust QTL identified by GWAS, including peak SNP, favorable allele, supporting genotypes, and linked gene content

**Table S5.** Genome annotation of genes located within LD-based QTL intervals associated with stripe rust resistance

**Table S6.** The list of high confidence QTLs supported by multiple stripe rust isolates or detected by both GWAS models and/or co-localized with previously designated Yr genes

**Table S7.** Watkins accessions carrying favorable alleles at high-confidence stripe rust resistance QTL supported by multiple isolates, GWAS models, or known *Yr* loci.

## Notes

### Competing Interest Statement

The authors have declared no competing interest.

